# Evolutionary history of the *α-Carbonic Anhydrase* (*αCA*) gene family in the phylum Arthropoda, with a focus on the copepod *Eurytemora affinis* species complex

**DOI:** 10.1101/2025.09.22.677836

**Authors:** Yifei Joye Zhou, Carol Eunmi Lee

## Abstract

The *α-Carbonic Anhydrase* (αCA) gene family encodes an abundant and ancient metalloenzyme present in all animals. Found in both eukaryotes and prokaryotes, members of this gene family play vital roles in ionic regulation, acid-base regulation, and respiration. However, little is known regarding the evolutionary history and patterns of molecular evolution of this gene family in the phylum Arthropoda. Through phylogenetic reconstruction and subcellular localization prediction, we discovered that arthropod αCA genes could be classified into three clades based on phylogenetic topology and predicted subcellular localization, as has been found in chordate αCA. Within the three distinct arthropod αCA clades, the Extracellular & Membrane-bound αCA clade exhibited the highest rates of evolution, variation in predicted subcellular localization, and number of homologs. Intriguingly, while we found that the majority of arthropod αCA homologs were classified in the Extracellular & Membrane-bound αCA clade, the majority of chordate αCA were within the Cytosolic αCA clade. This result indicates divergence in αCA expansion patterns between the two phyla. In addition, we found that αCA12, a paralog that showed salinity-associated signatures of selection in the copepod *Eurytemora affinis* species complex in previous studies, also shows signatures of positive selection (based on *dN/dS*) in two of the three sister species of the copepod *Eurytemora affinis* species complex (*E. carolleeae* and *E. gulfia*), compared to the general arthropod αCA background. Overall, our results indicate that the αCA gene family shows signatures of selection at the macroevolutionary scale, across phyla, and between closely related sister species.

**Significance statement:** *Alpha Carbonic Anhydrase* (αCA) is an essential enzyme that is particularly well-studied among certain metazoan lineages, such as chordates and poriferans, but not in arthropods. Here, we provide the first comprehensive analysis of the evolutionary history of αCA, and uncover the signatures of selection across phylum and between species. Through phylogenetic reconstruction, predicted subcellular localization, and *dN/dS*-based selection tests, our study shed light on how the αCA gene family has and continues to evolve in the largest animal phylum Arthropoda.

## Introduction

The *Alpha carbonic anhydrase* (αCA) gene family encodes metalloenzymes that are involved in a wide variety of essential biological processes, such as cellular respiration, ion transport, acid-base regulation, and photosynthesis (Badger et al., 1994; Esbaugh & Tufts, 2006; Gilmour & Perry, 2009; Henry & Cameron, 1983; Supuran, 2016). αCA enzymes catalyze the reversible reaction of H_2_O + CO_2_ ↔ HCO_3_^-^ + H^+^. The protons and bicarbonate (HCO_3_^-^) ions generated by this reaction are widely used in various major metabolic pathways, including ionic regulation (Lindskog, 1997; Lee et al., 2022). For instance, the export of αCA-generated protons and bicarbonate ions from the epithelial cells in exchange for Na^+^ and Cl^-^ is essential for many organisms living under low salinity conditions (Ali et al., 2015; Cardoso et al., 2019; Lee et al., 2022). In addition, functional co-option of αCA into biomineralization is well-characterized in certain metazoan taxa, including in mollusks, cnidarians, vertebrates, and sponges (Bertucci et al., 2013; Le Roy et al., 2014; Le Roy et al., 2016; Matt et al., 2022; Reibring et al, 2014). Further specialization of αCA occurs in the *Carbonic Anhydrase-Related -Proteins* (CARPs), a highly conserved but catalytically inactive clade within the αCA family that is suggested to have neural developmental functions (Aspatwar et al., 2010; Aspatwar et al., 2013; Aspatwar et al., 2022; Hirota et al., 2013; Taniuchi et al., 2002).

Among the families of carbonic anhydrases (α, β, γ, δ, ζ, η, θ, ι) that are genetically unrelated but have similar function, animals typically possess members of the αCA and βCA families (Aspatwar et al., 2022; Supuran et al., 2008; Supuran et al., 2016; Hassan et al., 2013; Jensen et al., 2019; DiMario et al., 2018). Also found in prokaryotes, plants, and fungi, the origin of αCA is proposed to have predated the emergence of animals (DiMario et al., 2017; Elleuche & Pöggeler, 2010). In addition, αCA exists as multiple paralogs within most animal genomes through multiple rounds of gene duplications, including an event dating to the Phanerozoic era (starting ∼541 mya) (Hewett-Emmett et al., 1984; Le Roy et al., 2014). Intriguingly, studies on vertebrate αCA have revealed the presence of diverse subcellular localization, including cytosolic, membrane-bound, transmembrane, mitochondrial, and extracellular localization (Hewett-Emmett, 2000; Hilvo et al., 2005; Aspatwar et al., 2022).

While αCA are ubiquitous and show intriguingly diverse patterns of localization in vertebrates, the αCA gene family in the phylum Arthropoda has long remained understudied. In fact, due to historically limited sequencing efforts, even the presence of αCA paralogs in certain arthropod subphyla had remained uncertain. Previous phylogenetic analyses of αCA have typically included only a few arthropod αCA sequences (mainly of hexapods), while focusing mainly on αCA in vertebrate or non-arthropod invertebrates (Banerjee & Deshpande, 2016; Le Roy et al., 2014). The only phylogenetic analysis on arthropod αCA focused on the genus *Daphnia* (Crustacea) (Culver & Morton 2015). Thus, the evolutionary history of arthropod αCA has remained unresolved.

Previous phylogenetic studies on vertebrate and non-arthropod invertebrate αCA have repeatedly found an ancient phylogenetic split between cytosolic/mitochondrial αCA and extracellular/membrane-bound αCA. This result suggests that the topology of the αCA phylogeny of animals is strongly associated with the patterns of αCA subcellular localization (Hewett-Emmett & Tashian, 1996; Le Roy et al., 2014; Jackson et al., 2007; Matt et al., 2022; Moya et al., 2008; Culver & Morton, 2015). However, the presence of such phylogenetic splits according to subcellular localization has not been explicitly tested for arthropod αCA.

Thus, a key question regards whether differences exist between patterns of αCA gene family expansions in arthropods *versus* chordates. Given that arthropods and chordates diverged ∼990 million years ago (Wang et al., 1999), arthropod αCA might show patterns of gene family expansion that differ from those of chordate αCA. Alternatively, given that αCA is an ancient gene family with an origin that likely predates the emergence of animal phyla, arthropod αCA clades might show similar patterns as chordate αCA, of ancient divergences associated with predicted subcellular localization of aCA enzymes (Le Roy et al., 2014; Jackson et al., 2007; Matt et al., 2022; Moya et al., 2008; Culver & Morton, 2015).

An especially intriguing feature of the αCA gene family is that members of this gene family are major targets of natural selection during salinity transitions in populations of the copepod *Eurytemora affinis* species complex. In particular, several αCA paralogs showed signatures of selection associated with salinity transitions in both wild populations and laboratory selection lines of *E. affinis* complex (Stern and Lee, 2020; Stern et al., 2022; Posavi et al., 2020; Lee, 2021; Lee et al., 2022). Repeatedly across different sibling species within the *E. affinis* species complex, αCA gene paralogs exhibited population genomic signatures of selection in response to salinity change and/or evolutionary shifts in gene expression (Lee, 2021). Specifically, out of the 13 αCA paralogs in the *E. affinis* complex genome, αCA1, αCA5, αCA9 and αCA12 showed signatures of selection in response to salinity change between saline and freshwater populations or during laboratory selection (Diaz et al., 2023; Du et al., 2025; Lee, 2021; Stern & Lee, 2020; Stern et al. 2022). In addition, certain αCA paralogs also showed evolutionary shifts in gene expression between saline and freshwater populations (Posavi et al., 2020). For instance, the paralog αCA9 of *E. carolleeae* displayed an evolutionary increase in gene expression under freshwater conditions in the freshwater-adapted population, relative to its saline ancestral population (Posavi et al., 2020). Given these previous results, we hypothesized that αCA paralogs of the *E. affinis* species complex would show signatures of positive selection (based on *dN/dS*), relative to other arthropod αCA paralogs.

Given the vast knowledge gap regarding the evolutionary history of arthropod αCA, this study aimed to address the hypotheses above, with some focus on αCA paralogs of the *E. affinis* complex. Specifically, the goals of this study were to (1) explore the evolutionary history of αCA paralogs across the phylum Arthropoda, (2) determine whether the clades classified based on predicted subcellular localization found in chordate αCA are also present in arthropod αCA, and (3) explore the phylogenetic signatures of positive selection and patterns of molecular evolution in αCA paralogs of the *E. affinis* species complex.

To accomplish these goals, we reconstructed a phylogeny of 294 αCA homologs from 31 arthropod species from all four arthropod subphyla (Chelicerata, Myriapoda, Crustacea, Hexapoda). We concurrently assigned predicted subcellular localization to each arthropod αCA homolog using DeepLoc 2.0 (Thumuluri et al., 2022). We then mapped the predicted subcellular localization of each arthropod αCA homolog onto the arthropod phylogeny, identifying the distinct αCA clades approximately based on predicted subcellular localization. The proportion of predicted subcellular localization within these distinct αCA clades were compared between arthropods and chordates to elucidate the potential evolutionary differences between αCA of the two phyla. Finally, for *E. affinis* complex αCA paralogs, we identified phylogenetic lineages and amino acid codons that showed signatures of positive selection based on *dN/dS*.

This study represents the first comprehensive analysis of the evolutionary history and distribution of predicted subcellular localization of the αCA gene family in the phylum Arthropoda. As a unique feature of this study, we included αCA sequences from all arthropod subphyla. Through our extensive taxon sampling, we were able to characterize patterns of gene family expansions in great detail, such as lineage-specific expansions of CARPs. We also characterized differential patterns of αCA paralog expansion, based on predicted subcellular localization, between chordate and arthropods αCA. Finally, our phylogenetic approach enabled us to investigate signatures of positive selection (based on *dN/dS*) within the protein structure of *E. affinis* complex αCA paralogs, relative to other arthropods, which previously showed population genomic signatures of selection. This signatures of selection within the αCA protein of *E. affinis* complex might be consistent with their extraordinary ability to cross salinity barriers while invading freshwater habitats, a feat rarely accomplished for invertebrates (Lee, 1999; Lee and Bell, 1999; Sługocki et al., 2021; Sukhikh et al., 2019).

## 2. Results and Discussion

### 2.1 Identification of αCA homologs across arthropod subphyla

From our phylogenetic analyses, we confirmed the existence of αCA genes in all 31 arthropod species included in our study (Supplementary Table S3). In addition, we found at least one CARP sequence in most of the arthropod genomes sampled (Supplementary Table S7). From the 31 arthropod species, we mined 353 distinct αCA sequences, including both partial and complete sequences, sampled across all 4 arthropod subphyla (Chelicerata – 8 species, Myriapoda – 4 species, Crustacea - 10 species, Hexapoda - 9 species).

Similar to previous vertebrate studies on αCA (Aspatwar et al., 2022), the number of αCA paralogs (non-CARP) varied greatly among arthropod taxa, with a mean of 9.19 ± 0.76 SE (Supplementary Material; Fig. S1, Supplementary Table S7). For example, in hexapods, non-CARP αCA paralog number varied from 4 in the gall adelgid *Adelges cooleyi* to 13 in the springtail *Folsomia candida*, with a mean value of 8.67± 0.87 SE. Crustaceans had a slightly higher number of non-CARP αCA paralogs (Mean = 11.20 ± 1.16 SE) than hexapods (Supplementary Material; Fig. S1 a&b; Supplementary Table S7). In addition, we found that the water flea, *Daphnia pulex*, had the highest number of non-CARP αCA paralogs (N = 19), followed by the Orange Rosary Millipede, *Helicorthomorpha holstii* (N = 18) (Supplementary Table S7). In contrast, CARP paralog numbers were lower and less variable among arthropod taxa, with a mean of 2.19 ± 0.31 SE. Most species harbored only 1-2 paralogs of CARPs (Supplementary Table S7).

In contrast to the low number of CARP paralogs in most arthropod taxa, we found elevated numbers in malacostracan crustaceans, which included 15 decapod species and 1 amphipod species (Supplementary Table S8; see Supplementary Methods). We mined CARP paralogs from 13 additional decapod and 1 additional amphipod genomes, beyond the 2 decapods used for subsequent analysis, to investigate the extent of the clade-specific CARP expansion (Supplementary Table S8; see Supplementary Methods). For the 15 decapod species included in this study (2 + 13 additional), the mean number of CARP paralogs was 9.00 ± 0.46 SE, much greater than the average CARP paralog number for arthropods (Mean = 2.19 ± 0.31 SE) (Supplementary Table S8). In addition, we found that the amphipod *Hyalella azteca*, another malacostracan outside the decapod clade, harbored 7 CARP paralogs in its genome. The mean number of CARP paralogs in the 15 decapod and one amphipod genomes (Mean = 8.81 ± 0.47) was significantly greater than the number in non-malacostracan crustaceans (Mean = 1.875 ± 0.30) (Wilcoxon rank-sum test; W = 128, *P* < 0.0005, Supplementary Material; Fig. S1d). Therefore, these results suggest that the CARP gene family expansion likely occurred prior to the split between Amphipoda and Decapoda, possibly affecting all members of the crustacean Class Malacostraca. Further taxon-sampling is required to fully support the hypothesis of malacostracan-specific CARP expansions, especially by including basal malacostracans, such as Syncaridans and Leptostracans, and additional crustacean taxa outside of malacostracans.

The expansion of CARP paralogs in Malacostraca is consistent with the dramatic rearrangement of neuroanatomy seen specifically in this clade (Strausfeld et al., 2020; Wolff & Strausfeld, 2015; Wolff et al., 2017). Given CARP’s function in neural development among chordates and its enriched expression in adult fly brain in *Drosophila* (Aspatwar et al., 2013; Aspatwar et al., 2014; Aspatwar et al., 2022), CARP’s expansion in the Malacostraca (particularly Decapoda) could potentially be linked to the derived neuroanatomy of this clade. Mushroom bodies of arthropod brains, the paired cognitive center responsible for learning and memory functions, is a conserved structure among arthropods that has undergone dramatic rearrangement in malacostracan crustaceans, especially in decapods (Strausfeld et al., 2020;

Wolff & Strausfeld, 2015; Wolff et al., 2017). Many members of Decapoda have lost various core features of the canonical mushroom body structure, including columnar lobes, while gaining various suborder-specific structures and novel arrangements of neuronal networks (Strausfeld et al., 2020). Within Decapoda, the neuroanatomy of brachyurans (true crabs) is especially derived, where even identification of neural centers with any resemblance to mushroom bodies has proven difficult (Strausfeld et al., 2020). In other malacostracans outside of decapods, stomatopoda possesses a canonical mushroom body structure that is reminiscent of that of hexapods, while Leptostraca has a highly diminished mushroom body (Kenning et al., 2013; Strausfeld et al., 2020; Wolff et al., 2017). All together, these results suggest that CARP paralogs might be playing a role in the extensive rearrangement of neuronal structure in malacostracans.

### 2.2 Phylogenetic reconstruction of arthropod αCA

To understand the evolutionary relationships among orthologs and paralogs of arthropod αCA, we performed phylogenetic analysis utilizing 294, from a total of 351 αCA sequences mined, distinct arthropod αCA homologs from 31 arthropod species, with 4 poriferan αCA paralogs as the outgroup.This phylogeny included paralogs from multiple species from all 4 arthropod subphyla (Chelicerata, Myriapoda, Crustacea, Hexapoda). We strived to include all αCA paralogs in the genome for each species.

Our Maximum-Likelihood phylogeny revealed the basal separation of all arthropod sequences into three major clades (Fig. 1a). To obtain further support for the topological separation of the three major clades, we also generated a UMAP projection of arthropod αCA homologs based on a protein language model, ESM-1b (Rives, Meier, Sercu et al., 2021). This analysis allowed us to determine whether the pattern of clustering based on structure-function properties captured by ESM-1b showed similar clustering as the sequence-based phylogenetic topology. This analysis was performed through an alignment-free sequence clustering method (Yeung et al., 2023) (see Methods and Materials). Interestingly, the resulting UMAP projection reconstructed the three distinct clades that we initially defined based on phylogenetic topology (Fig. 1b). Thus, our results support the basal separation of all arthropod sequences into 3 major clades through alignment-based phylogeny and non-alignment-based clustering.

**Figure 1.**
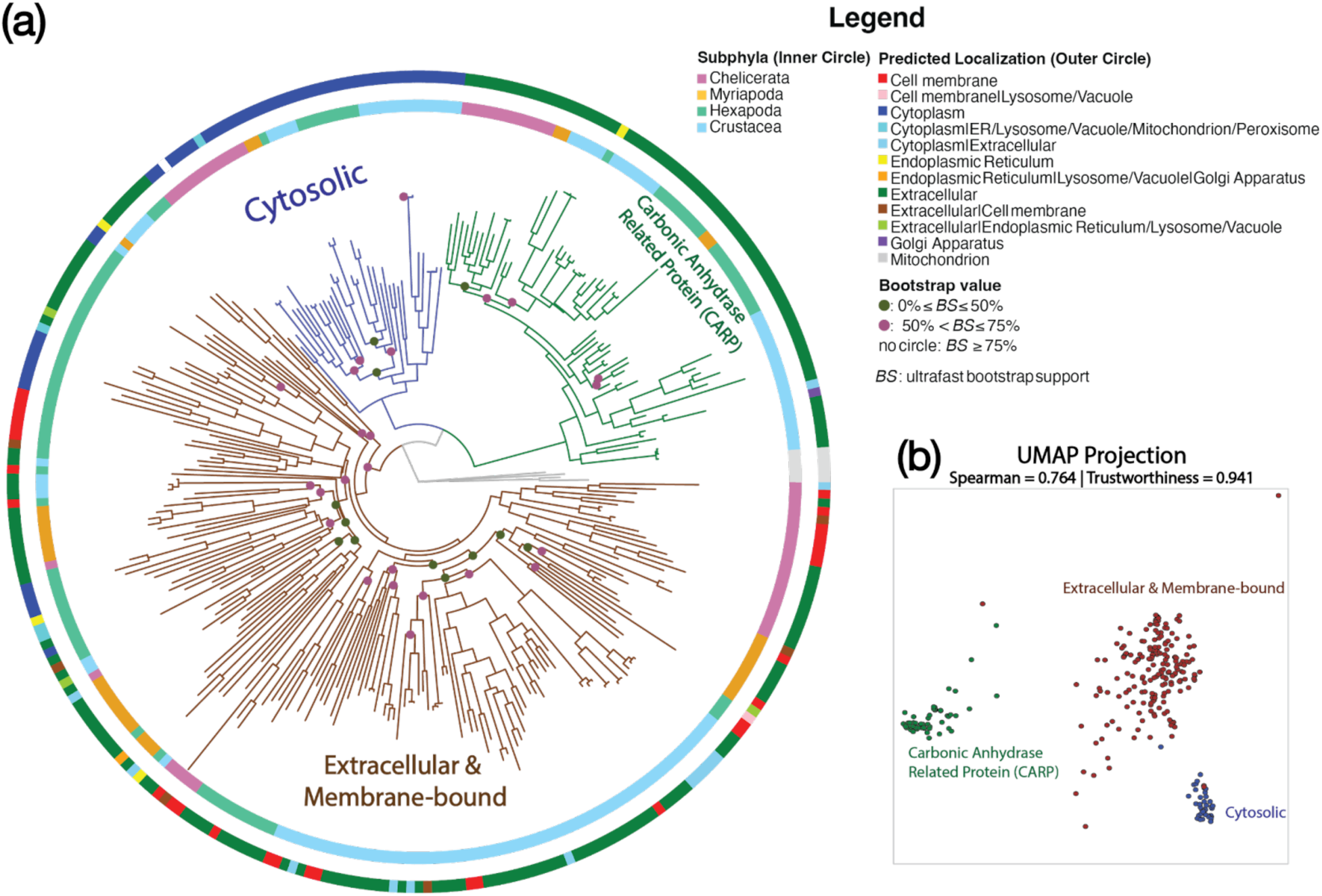
Arthropod αCA phylogeny with subcellular localization and UMAP projection. (**a**) Maximum Likelihood arthropod αCA phylogeny, including 294 distinct homologs sampled from all four arthropod subphyla, with Porifera αCA paralogs (grey branches) as the outgroup. The inner circle represents the arthropod subphyla. The outer circle represents the predicted subcellular localization of each paralog on this phylogeny based on DeepLoc 2.0 (Thumuluri et al., 2022). The three distinct clades of αCA (Extracellular & Membrane-bound, Cytosolic, and CARP) were designated based on phylogenetic topology and predicted subcellular localization, with phylogenetic topology taking precedence. The colored dots at the base of each node represents the bootstrap confidence levels. The nodes lacking dots have bootstrap values ≥ 75%. (**b**) A UMAP projection of 294 distinct arthropod αCA homologs, based on ESM-1b embedding (Yeung et al., 2023). The distance was calculated through TS-SS similarity, and the global and local accuracy of this clustering is shown through the Spearman rank correlation and trustworthiness values. Each dot represents a distinct αCA paralog, which was colored based on the associated αCA clade, determined from the phylogeny (Fig. 1a).

Given that previous phylogenetic studies on vertebrate and metazoan αCA have repeatedly found basal phylogenetic splits according to subcellular localization, we examined whether the 3 major clades of arthropod αCA phylogeny also showed divergences based on subcellular localization (Hewett-Emmett & Tashian, 1996; Le Roy et al., 2014; Jackson et al., 2007; Matt et al., 2022; Moya et al., 2008; Culver & Morton, 2015). We determined the predicted localization of each arthropod αCA paralog using DeepLoc 2.0 (Thumuluri et al., 2022). DeepLoc 2.0 uses a transformer-based deep-learning mode, trained on the UniProt dataset, to predict the subcellular localization of each query amino acid sequence (UniProt consortium, 2019; See Methods). To uncover the extent of congruence between the predicted subcellular localization and phylogenetic topology of arthropod αCA, we first mapped the predicted subcellular localization of each αCA homolog on to the phylogeny, and roughly assigned the main localization to each of the 3 main clades. We then determined the proportion of αCA sequences predicted to be in each localization within each phylogenetic clade (see Section 2.3 below).

The mapping of predicted subcellular localization revealed that the three major clades of arthropod αCA roughly differed in their predicted localizations (Fig. 1a, outer circle). The basal divergence among arthropod αCA sequences separated approximately into (1) Cytosolic (blue), (2) extracellular CARP (green), and (3) Extracellular & Membrane-bound (brown) clades, based on the most common localization within each clade (Fig. 1a). The cytosolic and extracellular

CARP clades formed sister clades within a larger clade. We also found a deep divergence between cytosolic and extracellular and membrane-bound αCA, consistent with previous αCA phylogenies (Hewett-Emmett & Tashian, 1996; Le Roy et al., 2014; Jackson et al., 2007; Matt et al., 2022; Moya et al., 2008; Culver & Morton, 2015). Therefore, we classify arthropod αCA approximately into three distinct clades based on localization and phylogenetic topology. However, note that not all members within these phylogenetic clades have the predicted subcellular localization of each respective clade (Fig. 1a, outer ring; see Section 2.3 below).

The topology of the arthropod αCA phylogeny (Fig. 1a; Supplementary Material; Fig. S2) was largely concordant with previous metazoan αCA phylogenies (Le Roy et al., 2014; Jackson et al., 2007; Matt et al., 2022; Moya et al., 2008). This congruence in overall topology of our arthropod αCA phylogeny (Fig. 1a) with previously reconstructed metazoan & vertebrate αCA phylogenies (Le Roy et al., 2014; Jackson et al., 2007; Matt et al., 2022; Moya et al., 2008) was not unexpected. αCA is an extremely ancient enzyme that is found across eukaryotes and even among different prokaryotes, especially in gram-negative bacteria (Capasso & Supuran, 2015; Hewett-Emmett & Tashian, 1996; Smith et al., 1999). Thus, the evolutionary origin of αCA likely preceded the emergence of eukaryotes. Therefore, we expected that the pattern of basal divergence to be largely consistent across any eukaryotic αCA phylogeny. In addition, the ancient evolutionary history of αCA might account for the low bootstrap support for the deep internal nodes and the long terminal branches in our arthropod αCA phylogeny (Fig. 1a).

We also found that αCA homologs from the four arthropod subphyla were present in all three arthropod αCA clades that we defined based on phylogenetic topology and predicted subcellular localization, consistent with previous αCA phylogenies that also found similar results in metazoans (Le Roy et al., 2014) (Figs. 2a & 2b, Supplementary Material; Figs. S2 & S3). This pattern also held true for paralogs of individual species, such as sibling species of the *E. affinis* species complex, which possess paralogs from all three clades (Fig. 2). These observations suggest that the most recent common ancestor of all arthropods possessed at least three αCA paralogs, with a paralog belonging to each of the three main αCA clades based on predicted subcellular localization.

**Figure 2.**
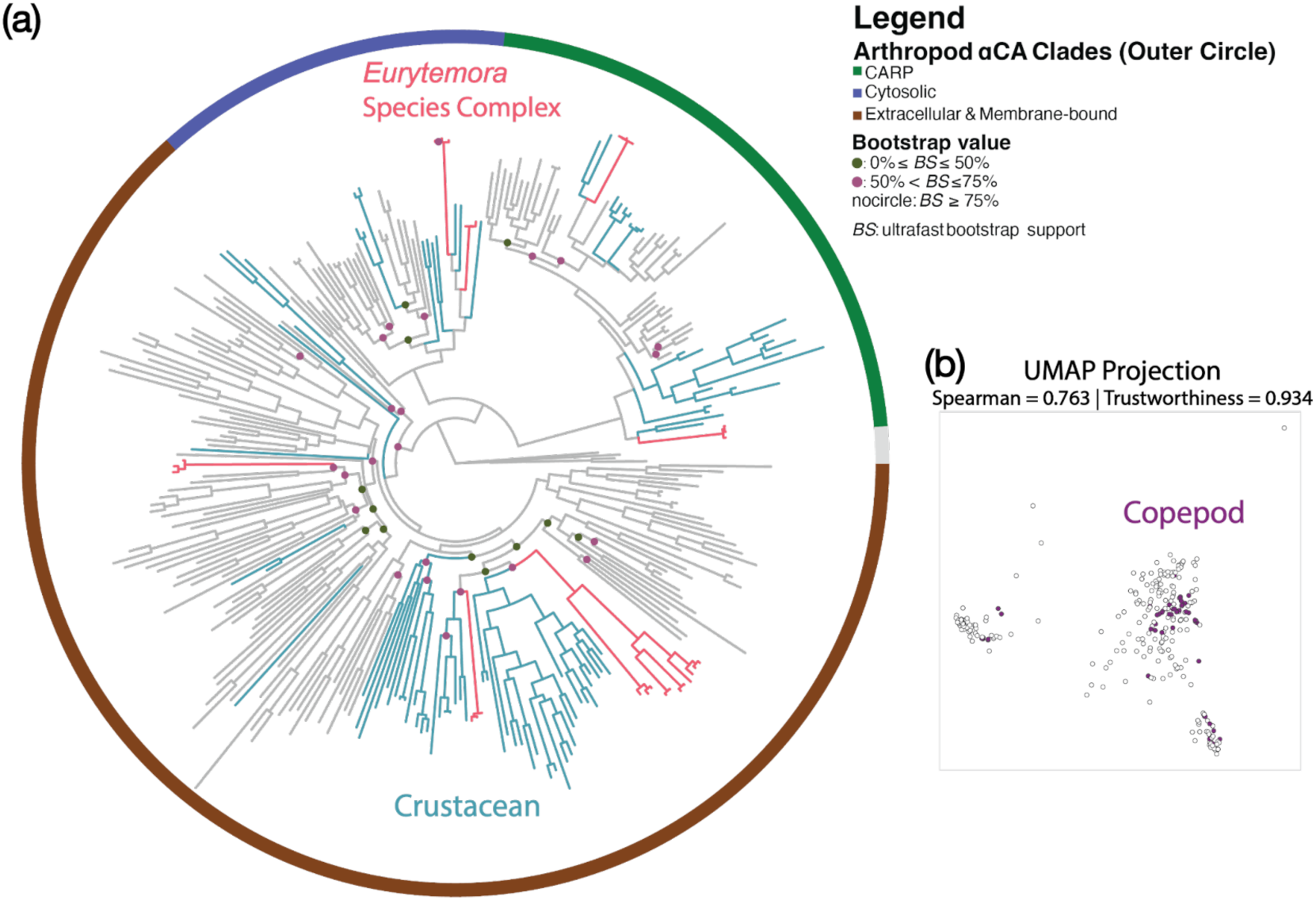
Arthropod αCA phylogeny showing crustacean branches (teal), especially *E. affinis* species complex (red), and UMAP projection. (**a**) Maximum likelihood arthropod αCA phylogeny as shown in Figure 1, but with the crustacean branches (teal) and *E. affinis* species complex (red) highlighted. This highlighting emphasizes the fact that all three clades based on predicted subcellular localization (Cytosolic, CARP, and Extracellular & Membrane-bound, outer circle, as in Fig. 1) occurs in crustaceans, including *E. affinis* complex.The colored dots at the base of each node represent bootstrap confidence levels. (**b**) A UMAP projection of 294 distinct arthropod αCA homologs based on ESM-1b embedding (Yeung et al., 2023) as seen in Figure 1. The distance is calculated through TS-SS similarity, and the global and local accuracy of this clustering is shown through a Spearman rank correlation and trustworthiness value. Each dot represents a distinct αCA homolog. Copepod αCA paralogs are colored in magenta.

Notably, our phylogenetic analysis revealed that arthropod αCA sequences grouped with chordate αCA sequences belonging to the same αCA clades based on predicted subcellular localization (Supplementary Material; Fig. S4). To investigate whether the arthropod αCA clades were homologous to the chordate αCA clades that were previously described (Hewett-Emmett & Tashian, 1996; Le Roy et al., 2014; Jackson et al., 2007), we reconstructed a reduced Maximum Likelihood phylogeny using a total of 198 αCA sequences from 10 arthropod species and 6 chordate species (See Supplementary Methods). The reconstructed phylogeny shows the clear grouping of arthropod and chordate αCA in each of the 3 αCA clades. Therefore, our findings suggest that arthropod and chordate αCA clades are homologous to each other. Furthermore, this result suggests that the most recent common ancestor of arthropods and chordates (Urbilateria) had at least three αCA paralogs, with a member in each of the three main αCA clades identified in this study (see previous paragraph).

While the presence of αCA in various prokaryotes supports the ancient origin of the αCA gene family, the evolutionary origin of the three distinct αCA clades, identified based on phylogenetic topology and predicted subcellular localization, remains unresolved. As mentioned above, our phylogenetic analysis between arthropod and chordate αCA suggests that the most recent common ancestor of arthropods and chordates (Urbilateria) had at least three αCA paralogs, with one in each of the distinct αCA clades (Fig. 2a, Supplementary Material; Fig. S4). Surprisingly, previous phylogenetic analyses have found that cnidarian αCA homologs also group with human αCA paralogs to form the three distinct αCA clades (Le Goff et al., 2016). This finding suggests that the 3 distinct αCA clades identified in this study have emerged before the divergence of Cnidaria and Bilateria. This ancient origin would suggest that the 3 distinct αCA clades are likely at least as ancient as the Hox/ParaHox homeodomain genes (Dunn et al., 2014; Ryan et al., 2010).

However, a question remains on the precise timing of origin of the three distinct αCA clades, particularly in relation to the basal branching animal phyla - Porifera (sponges) and Ctenophora (comb jellies). Poriferan αCA are often chosen as the outgroup of metazoan or vertebrate αCA because poriferan αCA homologs form a monophyletic group outside of the three distinct αCA clades (Jackson et al., 2007; Moya et al., 2012). This placement of poriferan αCA suggests that the three distinct αCA clades formed after the divergence between the most recent common ancestor of Porifera and Parahoxozoa (which includes Bilateria, Cnidaria, and Placozoa). In contrast, the metazoan phylogeny in Voigt et al. (2021) suggests that poriferan αCA are clustered with mammalian αCA into the three αCA clades. Therefore, whether the three distinct αCA clades, identified in this study, have originated before or after the split between poriferan and Parahoxozoa remains to be tested comprehensively. In addition, the presence of αCA genes in genomes of the basal phylum Ctenophora is unknown. Therefore, questions on the evolutionary origin of the three distinct αCA clades still require further exploration.

### 2.3 Diversity of localization and rates of evolution of the three arthropod αCA clades

As mentioned above (see Section 2.2), phylogenetic clade (Fig. 1a) and subcellular localization predicted by DeepLoc 2.0 were not always consistent. That is, the three αCA clades separated approximately according to predicted subcellular localization (Fig. 1a), but not completely. Overall, we found that αCA sequences that were classified phylogenetically into the extracellular CARP and Cytosolic αCA clades were largely consistent in their predicted subcellular localization, being consistent in 95.38% and 95% of the homologs respectively (Fig. 3a). However, the sequences that were classified phylogenetically into the Extracellular & Membrane-bound clade were consistent in their predicted subcellular localization only 77.25% (146 out of 189 sequences) of the time. That is, 22.75% (43 out of 189 sequences) of the sequences in the Extracellular & Membrane-bound clade (Fig. 1a) were not predicted to localize in either the extracellular space or on the cell membrane. Intriguingly, among the inconsistent homologs classified within the Extracellular & Membrane-bound clade, we also found multiple sequences with predicted cytosolic localization (Fig. 3a, rightmost bar in red). We also found that the extracellular & membrane-bound αCA clade contained the highest number of αCA sequences out of the three arthropod αCA clades (Fig. 1a & 3b).

**Figure 3.**
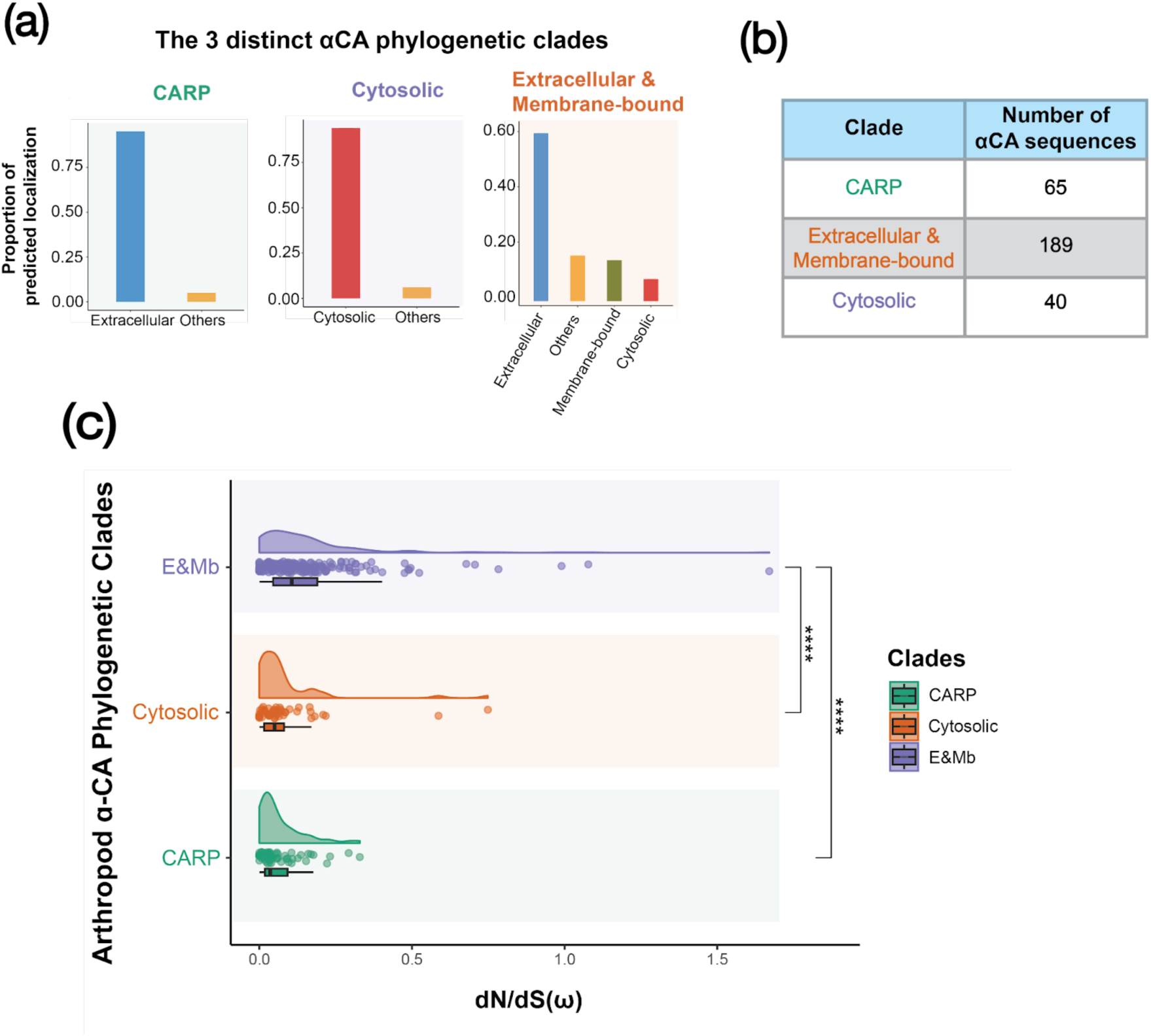
Proportion of predicted subcellular localization and rates of evolution in different αCA phylogenetic clades. (**a**) Proportion of αCA homologs with each predicted subcellular localization in different arthropod αCA phylogenetic clades. Any localizations other than Extracellular, Cytoplasm, and Cell Membrane are decoded as Others, including homologs with multiple potential localizations. CM is abbreviated for Cell Membrane localization. (**b**) Number of arthropod αCA homologs in each of the distinct αCA clades. (**c**) Rates of evolution (based on *dN/dS*) in different αCA clades, showing the fastest rates in the Extracellular & Membrane-bound αCA. Box and density plots showing the mean branch and node-specific ω value (*dN/dS*) of each arthropod αCA clade. E&Mb: mean ω = 0.16 ± 0.01 SE. Cytosolic: mean ω = 0.08 ± 0.02 SE. CARP: mean ω = 0.07 ± 0.01 SE. E&Mb: the Extracellular & Membrane-bound αCA clade. Pairwise comparison between CARP-E&Mb and Cytosolic-E&Mb using Wilcoxon rank-sum test; ns = not significant; * = p<0.05; ** = p<0.01; *** = p<0.001; **** = p<0.0001

Given the high variation in numbers of sequences belonging to each arthropod αCA clade (Fig. 3b), we hypothesized that evolutionary rates would vary among the three clades. In particular, based on the high numbers of αCA homologs in the Extracellular & Membrane-bound clade (Fig. 3b), we predicted that signatures of selection would be most prominent in this clade. Thus, we explored signatures of selection (in terms of *dN/dS* = ω, using FitMG94 offered in the HyPhy software suit) along the phylogenetic branches of the arthropod αCA phylogeny (Muse & Gaut, 1994; Pond et al., 2005). For this analysis, we conducted stringent trimming of the sequence alignment to reduce saturation and false positives, keeping only codons with conservation levels that were at least 50% in the alignments (see Methods and Materials). Our results did indeed reveal that Extracellular & Membrane-bound αCA had significantly higher branch and node ω values (*dN/dS*), relative to both the Cytosolic and CARP αCA clades (Wilcoxon rank-sum test; W = 3146.0 and 3492.5, respectively, *P* < 0.0001 for both comparisons) (Fig. 3c). These results were consistent with positive selection driving the diversification of αCA homologs in the Extracellular & Membrane-bound αCA clades.

Intriguingly, we found that the higher signatures of selection in the Extracellular & Membrane-bound αCA clade in arthropods (previous paragraph) also occurred in chordates (Supplementary Material; Fig. S5). We examined signatures of selection (based on *dN/dS)* for the three different chordate αCA clades. As in the case for arthropod αCA clades (Fig. 3c), we found that chordate Extracellular & Membrane-bound αCA had significantly higher branch and node ω values (*dN/dS*) than Cytosolic and CARP αCA (Supplementary Material; Fig. S5). A similar result, with extracellular αCA showing higher rates of evolution (based on *dN/dS*) than the intracellular αCA, was also seen in a previous study conducted on vertebrate αCA (McDevitt & Lambert, 2011).

Based on our results (Fig. 3c, Supplementary Material; Fig. S5), the rapid evolutionary rates in the Extracellular & Membrane-bound αCA clade appear to predate the formation of the arthropod and chordate clades. However, the causes of the elevated evolutionary rates in the Extracellular & Membrane-bound αCA clades remain unresolved. It is worth noting that extracellular proteins are generally known to demonstrate higher than average rates of evolution than intracellular proteins, regardless of the functions of the genes (Julenius & Pederson, 2006). Future research is needed to decipher the biological mechanisms leading to the elevated evolutionary rates seen in the Extracellular & Membrane-bound αCA clade.

### 2.4 Differential αCA clade expansions in arthropods versus chordates

For the 3 distinct αCA clades present in both arthropods and chordates (Fig. 1), we found stark differences in patterns of αCA gene family expansions. We found a high proportion of αCA homologs belonging to the Extracellular & Membrane-bound clade in arthropod species (64.3%) and a low proportion belonging to the Cytosolic clade (13.6%). In contrast, in chordates, relative to arthropods, we found a relative enrichment in αCA homologs belonging to the Cytosolic clade (37.5%), but a relative reduction of αCA homologs in the Extracellular & Membrane-bound clade (44.4%) (Fig. 4a). In a pairwise comparison, arthropod species showed significantly higher proportions of homologs in the Extracellular & Membrane-bound clade than chordates (Wilcoxon rank-sum test; W = 96.5, *P* = 0.0103) (Fig. 4b, right graph), whereas chordate species showed significantly higher proportions of Cytosolic αCA paralogs than arthropod species (Wilcoxon rank-sum test; W = 2.5, *P* = 0.00115) (Fig. 4b, left graph).

**Figure 4.**
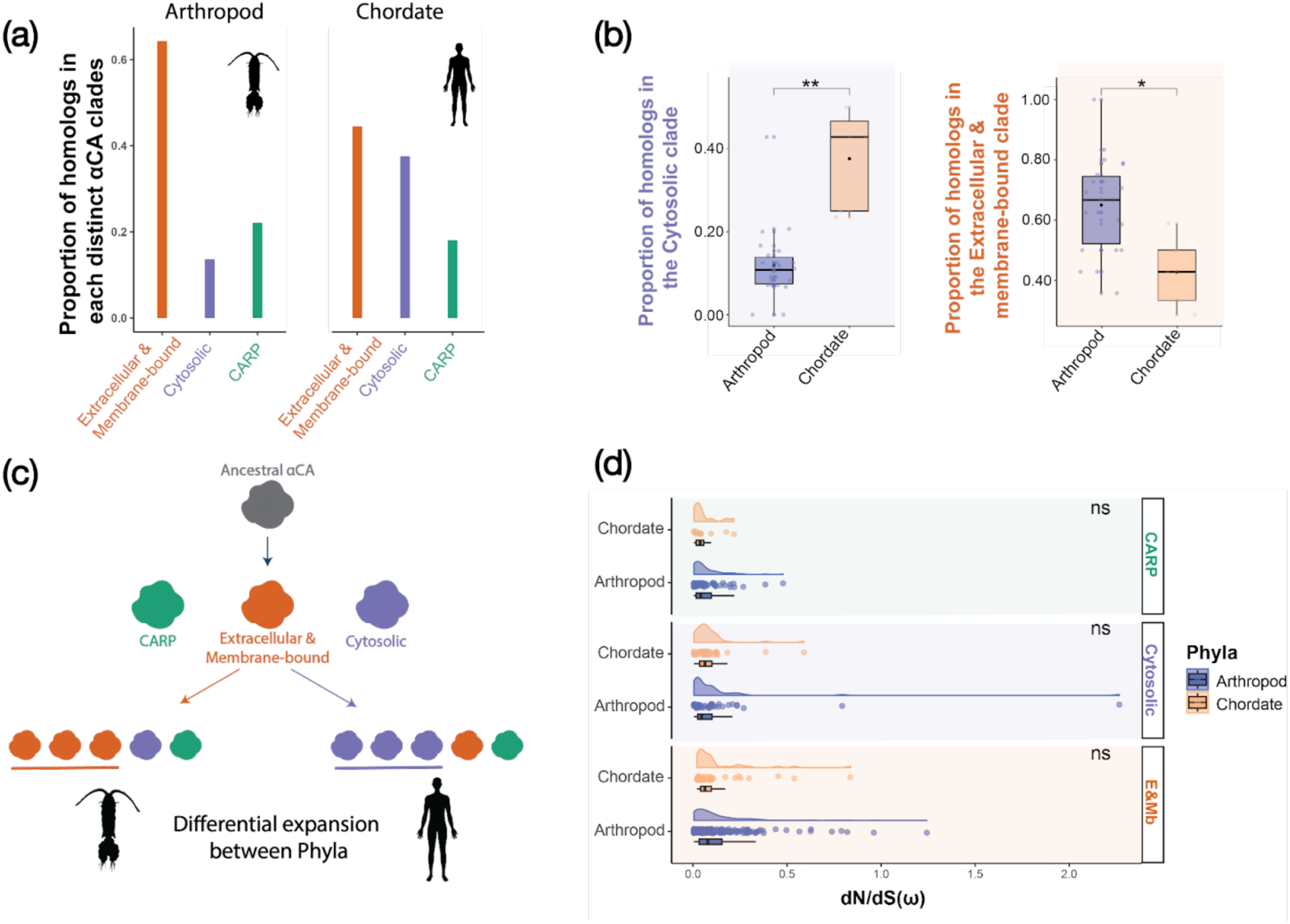
Differential patterns of αCA gene family expansion patterns in arthropods versus chordates. (**a)** Proportion of αCA homologs (orthologs and paralogs) within the 3 distinct αCA clade classified according to phylogenetic topology for arthropod αCA (left) and chordate αCA homologs (right). (**b**) Mean proportion among species of homologs that are classified in the Cytosolic clade (left) or classified in the Extracellular & Membrane-bound clade (right) between arthropod and chordate species. Wilcoxon rank-sum test; ns = not significant; * = *P* < 0.05; ** = *P* < 0.01. Cytosolic: Arthropod mean = 0.12 ± 0.06 SE; Chordate mean = 0.38 ± 0.06 SE. E&Mb: Arthropod mean = 0.65 ± 0.03 SE; Chordate mean = 0.42 ± 0.05 SE. (**c**) Depiction of differential αCA expansion patterns in arthropods versus chordates. (**d**) Rates of evolution (based on *dN/dS*) in different αCA clades for both arthropods and chordates. Box and density plots show the branch and node ω value (*dN/dS*) calculated based on FitMG94 implemented in the HyPhy software suite of each type of αCA clade in arthropods and chordates (Muse & Gaut, 1994; Pond et al., 2005). E&Mb refers to the Extracellular & Membrane-bound αCA clade. E&Mb: Arthropod: mean ω = 0.12 ± 0.01 SE, Chordate: mean ω = 0.05 ± 0.02 SE. Cytosolic: Arthropod: mean ω = 0.13 ± 0.05 SE; Chordate: mean ω = 0.08 ± 0.02 SE. CARP: Arthropod: mean ω = 0.06 ± 0.01 SE; Chordate: mean ω = 0.05 ± 0.02 SE.

We found similar patterns of enrichment when we compared αCA homologs classified only by predicted subcellular localization using DeepLoc 2.0, rather than with phylogenetic clades (Thumuluri et al., 2022) (Supplementary Material; Fig. S7). Similar to the patterns found above (previous paragraph), we found an enrichment of αCA homologs with predicted Extracellular & Membrane-bound localization in arthropods and an enrichment of αCA homologs with predicted Cytosolic localization in chordates. We further found that arthropod species showed significantly higher proportions of homologs with predicted extracellular localization than chordates in a pairwise comparison (Wilcoxon rank-sum test; W =105, *P* = 0.00198) (Supplementary Material; Fig. S7).

Although more sampling of animal phyla would be needed to determine the timing of origins of the subcellular localization clades, the phylum-specific patterns αCA family expansions (Fig. 4c) is particularly intriguing, and their evolutionary causes and functional consequences warrant further investigation.

Given the divergent patterns of expansions of αCA clades between chordates and arthropods, we hypothesized that evolutionary rates would also differ between arthropod versus chordate αCA lineages within the Extracellular & Membrane-bound or Cytosolic clades. We specifically predicted that rates of evolution would be greater in the clade that experienced greater expansions (Fig. 4a-c). Therefore, we expected that homologs from the Extracellular & Membrane-bound clade would evolve relatively faster in arthropods, whereas homologs from the cytosolic clade would evolve relatively faster in chordates. The reasoning is that after gene duplications, we expect the rate of evolution of the duplicated genes to rise due to relaxation of purifying selection and/or positive selection (Ohno, 2013; Panchy et al., 2016; Rispe et al., 2008; Scannell & Wolfe, 2008). Under the process of neofunctionalization or subfunctionalization, the duplicated αCA genes that are retained in the chordate or arthropod genomes might gain a different or more specialized function compared to the ancestral paralog (Ohno, 2013). This divergence in function might lead to divergent directional selection between the homologs, keeping the rate of evolution (*dN/dS*) high even after the initial relaxation of purifying selection.

To test the hypothesis that greater gene family expansions would correspond to greater rates of evolution, we compared signatures of selection between αCA clades of arthropods and chordates. Specifically, we calculated the branch and node-specific ω values (*dN/dS*) through FitMG94, as implemented in the HyPhy software suite for each αCA clade (Muse & Gaut, 1994; Pond et al., 2005). However, we found no significant differences in mean ω value in pairwise comparisons between arthropod and chordate αCA for each subcellular localization clade (Fig. 4d) (Wilcoxon rank-sum test; arthropod vs chordate for CARP: W = 452, *P* = 0.68. Cytosolic: W = 920, *P* = 0.62, extracellular and membrane-bound: W = 4183, *P* = 0.96). A potential explanation for this result is that long evolutionary time has passed since the phyla-specific expansion of αCA clades. As evolutionary time increases, the observed relaxation of purifying selection in recently duplicated homologs diminishes. In addition, the ω values (*dN/dS*) would become eroded through the accumulation of multiple DNA mutations at each nucleotide site, especially when the proportion of the gene that is under strong purifying selection is small (Shapiro & Elm, 2009). These factors could lead to *dN/dS* being inefficient at detecting ancestral positive selection events.

### 2.5 Signatures of positive selection at αCA paralogs of the *Eurytemora affinis* species complex

We were particularly interested in testing whether *E. affinis* complex αCA paralogs that had previously shown salinity-associated signatures of positive selection during contemporary shifts in habitat salinity (Lee, 2021; Stern and Lee, 2020; Stern et al., 2022; Posavi et al. 2020) also showed more ancient signatures of positive selection relative to other arthropod αCA. While the previous studies detected signatures of selection based on allele frequency shifts (e.g., *F*_ST_, haplotype diversity) during recent salinity change (over decades), the analysis here explored selection signatures using *dN/dS* between species separated by millions of years. To be more specific, we used aBSREL in HyPhy to detect signatures of positive selection along phylogenetic branches (Smith et al., 2015). We had strong rationale for making this comparison, given our previous studies that found that specific ion transport-related genes are under selection in response to salinity change on both contemporary and ancient time scales (Du et al. 2024; Du et al. 2025).

To identify signatures of selection using HyPhy, we generated alignments of arthropod αCA using conserved sequences to reduce phylogenetic noise and avoid obtaining false positive signatures of selection. To identify the conserved codon sites, we implemented a stringent trimming process, where only codons with conservation level of ≥60% were retained (see Methods and Materials). We selected all αCA paralogs of the *E. affinis* species complex as the foreground for this analysis. For the three αCA clades, we ran the trimming and the positive selection analyses separately. For instance, all Extracellular & Membrane-bound αCA of the *E. affinis* complex was compared only to other arthropod Extracellular & Membrane-bound αCA during both the trimming process and selection analyses.

In this analysis, we found only one phylogenetic position (node/branch) with signatures of positive selection across all αCA paralogs of the *E. affinis* complex, localized at αCA12 (Fig. 5a). Specifically, the branch leading to the clade grouping αCA12 of *E.carolleeae* (Atlantic clade) with *E. gulfia* (Gulf clade), and diverging from αCA12 of the basal *E. affinis* proper (Europe clade), showed signatures of positive selection (Fig. 5a, red branch; aBSREL; ω2 = 1443; Bonferroni-corrected *P* = 0.003). Intriguingly, αCA12 is also a paralog within the Extracellular & Membrane-bound αCA clade that has shown population genomic signatures of selection (allele frequency shifts) associated with shifts in habitat salinity in previous studies for both *E. carolleeae* and *E. affinis* proper (Table 1) (Stern & Lee, 2020; Stern et al. 2022; Diaz et al., 2023). These results support the hypothesis that αCA of the *E. affinis* species complex that showed contemporary signatures of selection associated with salinity shifts in previous studies also show ancient signatures of positive selection in comparison with other arthropod αCA paralogs.

**Figure 5.**
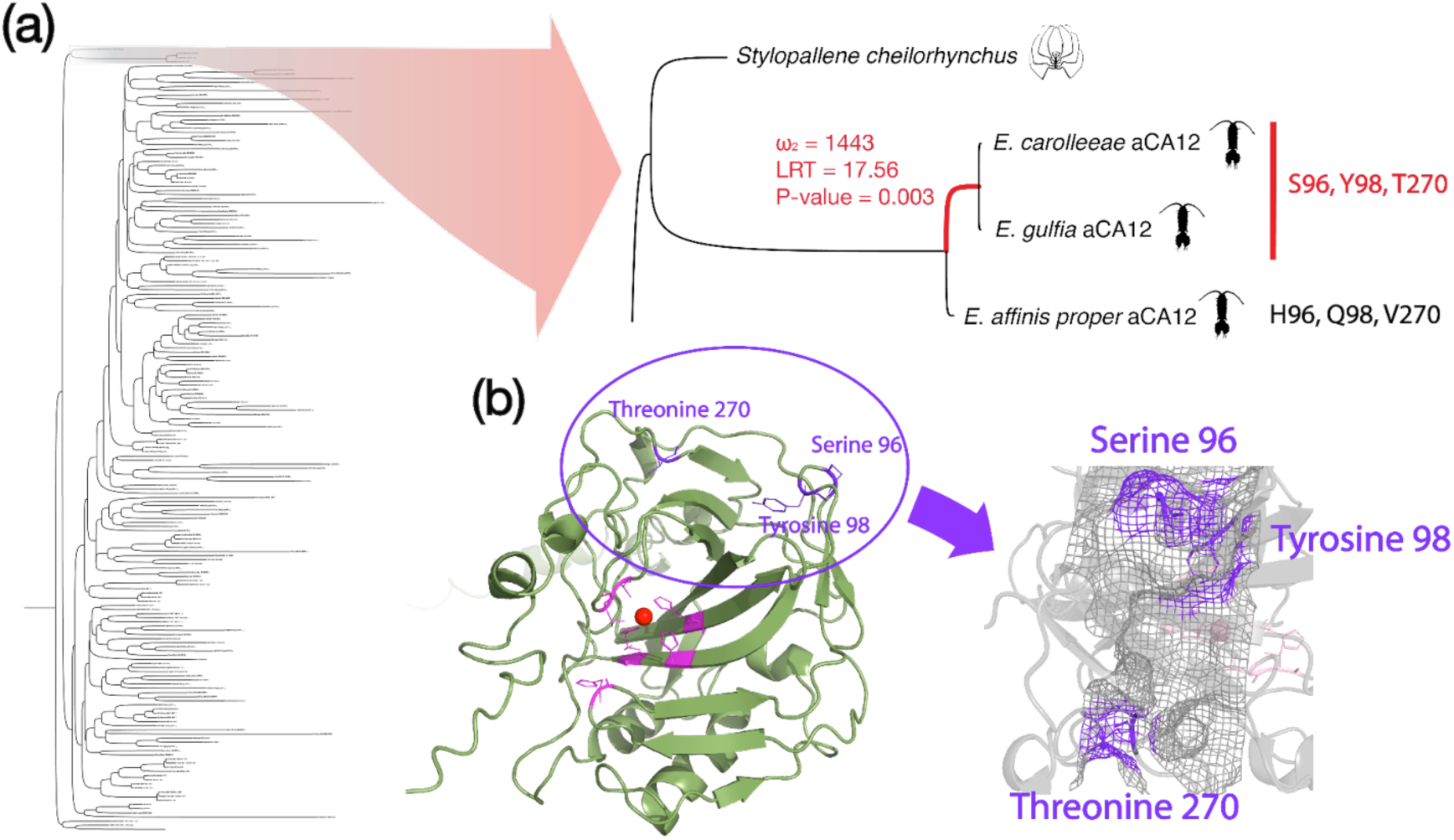
Signatures of positive selection in the arthropod Extracellular & Membrane-bound αCA phylogeny and at amino acid sites in the αCA12 protein model of *E. carolleeae*. (**a**) Arthropod Extracellular & Membrane-bound αCA phylogeny (left), highlighting the branch showing signatures of positive selection (right, red branch). aBSREL; ω2 = 1443; LRT = 17.56; Bonferroni-corrected *P* = 0.003. The amino acid changes at the sites with signatures of positive selection are shown in red font, in comparison to the ancestral state shown in black. These amino acid changes correspond to the same codons under selection in (b) shown in purple font. (**b**) Protein model of αCA12 of *Eurytemora carolleeae*, generated using Alphafold3. The nucleotide sites colored in magenta (left model) signifies the locations of the active pockets that catalyze the reverse hydration of CO_2_ (Hassan et al., 2013).The red sphere represents the Zn^2+^ ion. The 3 residues in the model marked in purple are the sites that are contributing to the signal of positive selection detected with BUSTED and MEME in HyPhy (Murrel et al., 2012; Murrel et al., 2015). The model on the right shows a magnified view of the residues contributing to the signals of selection. The numbers next to the residues represent the positions of the residues on the protein.

**Table 1.** Predicted localization of aCA paralogs of the copepod *Eurytemora affinis* complex that showed evolutionary responses to salinity change in prior studies. Shown are *E. affinis* complex αCA paralogs with their phylogenetic clade, predicted subcellular localization, population genomic signatures of selection in response to contemporary salinity change (Stern and Lee, 2020; Stern et al., 2022; Diaz et al., 2023), and evolutionary shifts in expression in response to salinity change (Posavi et al., 2020). αCA3 and αCA6 were omitted from the analysis as they were partial genes.

| #Paralogs | Phylogenetic Clade (Fig. 1a) | Predicted Subcellular Localization | Population Genomic Signatures of Selection during Contemporary Salinity Change | Evolution of Gene Expression between Saline vs. Freshwater populations<sup>d</sup> |
| --- | --- | --- | --- | --- |
| αCA1 | Extracellular & Membrane-bound | Cytoplasm/Extracellular | Yes<sup>a</sup> | N.S. |
| αCA2 | CARP | Extracellular | Yes<sup>c</sup> | N.S. |
| αCA3 | - | - | - | - |
| αCA4 | CARP | Extracellular | No | N.S. |
| αCA5 | Cytosolic | Cytoplasm | Yes<sup>a</sup> | N.S. |
| αCA6 | - | - | - | - |
| αCA7 | Cytosolic | Cytoplasm | Yes<sup>c</sup> | Up in saline population |
| αCA8 | Extracellular & Membrane-bound | Extracellular | No | Up in saline population |
| αCA9 | Extracellular & Membrane-bound | Cytoplasm/Extracellular | Yes<sup>c</sup> | Up in freshwater population |
| αCA10 | Extracellular & Membrane-bound | Cytoplasm/Extracellular | No | N.S. |
| αCA11 | Extracellular & Membrane-bound | Extracellular | No | Up in saline population |
| αCA12 | Extracellular & Membrane-bound | Extracellular | Yes<sup>a,b,c</sup> | Up in saline population |
| αCA14 | Extracellular & Membrane-bound | Extracellular | No | Up in saline population |

<sup>a</sup>Stern and Lee, 2020
<sup>b</sup>Stern et al. 2022
<sup>c</sup>Diaz et al. 2023
<sup>d</sup>Posavi et al. 2020

To understand the specific amino acid codon sites that showed signatures of positive selection in αCA12 (Fig. 5), we used BUSTED, with its Evidence Ratio (ER) value as a proxy, and MEME in the software HyPhy (Murrel et al., 2012; Murrel et al., 2015). We have strong rationale to focus on αCA12 as it is the only αCA paralog that showed population genomic signatures of positive selection in three sibling species (*E. carolleeae*, *E. gulfia*, *E. affinis* proper) within the *E. affinis* species complex, either in response to contemporary salinity change or during laboratory selection (Table 1; Stern & Lee, 2020; Stern et al. 2022; Diaz et al., 2023). In addition, αCA12 also shows evolution of increased expression in the freshwater population of *E. carolleeae* under saltwater conditions relative to its saline ancestor (Table 1; Posavi et al., 2020).

The evidence ratio (ER) is a likelihood ratio between the alternative model (presence of positive selection at a given amino acid codon site) and the null model (no positive selection at a given amino acid codon site), providing descriptive information on whether a site might have evolved under positive selection. In this analysis, we used αCA12 paralogs of *E. carolleeae* and *E. gulfia* and their branching node as the foreground against the rest of the arthropod Extracellular & Membrane-bound αCA paralogs as the background.

As a result, we found codon-specific signatures of positive selection on codon 21 of the edited alignment through MEME (α = 3.78, β^+^ = 128.12, *P* = 0.01) (See Github repository). In addition, our BUSTED analysis revealed gene-wide signatures of positive selection at the αCA12 paralogs of *E. carolleeae* and *E. gulfia*, compared to the background (*P* = 0.0035), with codons 21,22, and 79 of the edited alignment potentially contributing to the selection signal (ER = 46.8, 16.3, 38.8, respectively) (See Github repository). These αCA12 codons (21,22, 79 on the edited alignment) under possible signatures of positive selection correspond to amino acid positions Ser96, Tyr98, and Thr270→ [are these the derived or ancestral states?], respectively, on the complete *E. carolleeae* αCA12 sequence (Fig. 5b). Results from both BUSTED and MEME converged on codon 21 (Ser96) as a target of positive selection when compared to the general arthropod αCA background. However, identifying codons under selection using BUSTED and ER values to analyze branches defined by aBSREL is an exploratory analysis intended to generate hypotheses. Further experimental testing and statistical analyses of these amino acid sites are required to identify the functions of these substitutions and whether they contribute to adaptive evolution of αCA12.--> so what are the amino acid substitutions?

To determine the physical positions of the amino acid codon sites with signatures of positive selection, we mapped these sites onto a predicted protein model of *E. carolleeae* αCA12, generated using AlphaFold3 (Abramson et al., 2024). In comparison to the ancestral sequence of *E. affinis* proper αCA12, we found that these codons in *E. carolleeae* and *E. gulfia* have all gained missense mutations, shifting from His96 to Ser96, Glu98 to Tyr98, and Val270 to Thr270 (Fig. 5a). We found that the 3 codons that contributed to the signals of gene-wide signatures of positive selection (Ser96, Tyr98, Thr270) (Fig. 5a, red font, Fig. 5b, purple font) were positioned on the surface of the protein, at a large distance from the proton shuttle and the entrance to the active pocket (Fig. 5b, magenta shading).

While the amino acid sites with signatures of positive selection are localized on the surface of the αCA12 protein model (Fig. 5b), far from the protein active site, some of the substitutions might impact the functions of the protein. For the histidine to serine mutation at position 96 (His96Ser), despite the massive reduction in size, both residues are polar and might not affect the overall structure of the protein (Betts & Russell, 2003). In contrast, the substitution from glutamate to tyrosine at position 98 (Glu98Tyr) involves replacing a negatively charged residue with a bulky, aromatic residue that is hydrophobic (Betts & Russell, 2003). As a result, this substitution might incur a structural effect on the αCA12 protein. Lastly, the valine to threonine substitution at position 270 (Val270Thr) entails replacing a hydrophobic residue with a small polar residue (Betts & Russell, 2003). Both serine and threonine are known to be common targets of multiple post-translational modification, such as phosphorylation and O-linked glycosylation (Betts & Russell, 2003; Bradley, 2022; Di Fiore et al., 2020). Perhaps these mutations could initiate novel regulatory mechanisms in *E. carolleeae* and *E. gulfia* αCA12.

Furthermore, It is intriguing that the node that leads to αCA12 of *E. carolleeae* and *E. gulfia* showed signatures of selection, while αCA12 of *E. affinis* proper did not. Based on a phylogeny reconstructed from whole genome sequences, *E. affinis* proper (Europe clade) is known to be basally branching relative to *E. carolleeae* and *E. gulfia* (Du et al., 2025), consistent with the αCA12 phylogeny (Fig. 5a). The signatures of selection at αCA12 might be consistent with functional evolution related to habitat expansions in the derived clades of *E. affinis* complex. Populations of *E. carolleeae* and *E. gulfia* are known to more frequently invade novel salinities than populations of *E. affinis* proper, with *E. carolleeae* having the highest number of freshwater colonizing populations among sibling species of the *E. affinis* species complex (Du et al., 2024; Sługocki et al., 2021; Sukhikh et al., 2013; Sukhikh et al., 2019). However, the specific functions of these amino acid sites under selection in *E. carolleeae* and *E. gulfia,* relative to *E. affinis* proper, remains to be investigated.

On the other hand, another paralog, αCA9, which had previously been implicated in freshwater adaptation in the *E. affinis* complex, showed no signatures of selection in our analyses (Supplementary Material; Fig. S10). This paralog had previously exhibited the evolution of increased expression in the freshwater population of *E. carolleeae* under freshwater conditions relative to its saline ancestor (Table 1; Posavi et al., 2020). Here, we found a lack of signatures of positive selection (lack of elevated *dN/dS*) for all codon sites included in our analysis of αCA9 paralogs of the *E. affinis* species complex, compared to the arthropod background. Therefore, αCA9 likely did not undergo significant amounts of sequence evolution in functional codon sites in *E. affinis* complex. Previous studies had found the lack of population genomic signatures of selection at αCA9 during saline to freshwater invasions in wild populations of *E. carolleeae* and *E. gulfia* (Stern and Lee, 2020). Although, signatures of selection in non-coding sequence near αCA9 were found in Baltic Sea populations of *E. affinis* proper along a salinity gradient (Diaz et al., 2023). The mechanisms of functional specialization of *E. affinis* complex αCA9 paralogs might arise from regulatory evolution alone and not from selection on amino acid sequences.

### 2.6 Signatures of positive selection among non-*Eurytemora* arthropod αCA

Outside of αCA paralogs of the *E. affinis* species complex, we also explored other arthropod αCA homologs for signatures of positive selection using aBSREL in HyPhy (Smith et al., 2015). We tested all arthropod αCA branches for signatures of positive selection as an exploratory analysis.

Intriguingly, we found branch-level signatures of positive selection in the terminal branch leading to a CARP paralog of the Chinese white shrimp *Penaeus chinensis* (Crustacea:Decapoda) (Penaeus_chinensis_XM_047624972_1_CARP), among the arthropod CARP sequences (Supplementary Material; Fig. S11). This result is particularly intriguing because we found evidence for a malacostracan-specific CARP expansion within our previous analyses (See Section 2.1). This result suggests that the new CARP paralogs, formed from malacostracan-specific CARP expansion, have higher rates of evolution when compared to the general arthropod CARP background, potentially due to the neofunctionalization and subfunctionalization of duplicated paralogs (Conant & Wolfe, 2008; Innan & Kondrashov, 2010; Kondrashov et al., 2002).

## 3. Conclusions

In summary, this study represents the most comprehensive phylogenetic analysis of arthropod αCA to date. By investigating the proportion of αCA homologs in the 3 distinct αCA clades, based on predicted subcellular localization, we provide evidence for divergent patterns of αCA expansion between chordate and arthropods. Our results bolster the hypothesis that the evolutionary origin of the 3 distinct αCA clades likely predated the divergence between bilateria and Cnidaria/Placozoa. Intriguingly, we also demonstrate that αCA12 of the *E. affinis* species complex, which showed contemporary salinity-associated signatures of positive selection in previous studies, also show ancient signatures of positive selection in comparison to other arthropod αCA paralogs. These results are consistent with αCA genes contributing to salinity adaptation in *E. affinis* complex on multiple time scales. This study offers the first foray into patterns of evolution of the expansive and rapidly evolving αCA gene family in the largest animal phylum Arthropoda.

## 4. Materials and Methods

### 4.1 Taxon Sampling, Gene Mining, and Paralog Identification

Nucleotide and amino acid sequences of 351 Alpha Carbonic Anhydrase genes (αCA) were mined from a total of 31 arthropod species from available genome and transcriptome databases (Supplementary Tables S1 & S3). Of these sequences, 294 arthropod αCA sequences were retained after quality control filtering and used for subsequent phylogenetic analyses (see below). In addition, 4 coding sequences of αCA were identified from the sponge *Amphimedon queenslandica* (Phylum Porifera), the outgroup taxon. Sequences were screened for pseudogenes, which were removed from the initial dataset, resulting in the 351 αCA paralog sequences used for this study. The phylogenetic analysis (see below) included αCA sequences from all 4 arthropod subphyla, including from 8 species of Chelicerata, 4 species of Myriapoda, 10 species of Crustacea, and 9 species of Hexapoda (Supplementary Material; Table S1). To explore signatures of natural selection between chordate and arthropod αCA clades based on *dN/dS* (the ratio of nonsynonymous (*dN*) and synonymous (*dS*) substitution rates) using the software package HyPhy (Pond et al. 2005, see below), an additional 72 αCA homologs (orthologs and paralogs) were mined for 6 chordate species (lancelet *Branchiostoma lanceolatum*, tunicate *Ciona intestinalis*, zebrafish *Danio rerio*, *Homo sapiens*, frog *Xenopus tropicalis*, and red junglefowl *Gallus gallus*) (Supplementary Material; Table S2).

The majority of these putative arthropod αCA sequences were mined from the NCBI database (Sayers et al. 2024) (Supplementary Material; Table S1). For hexapods and crustaceans, genomes with good underlying quality (BUSCO score > 95% and/or contig N50 > 1Mb) were pre-selected, and BLAST searches (tblastn) were conducted on each of these genomes specifically (Altschul et al., 1990; Camacho et al., 2009; Gertz et al., 2006; Johnson et al. 2008; Simão et al., 2015). For the sibling species within the *E. affinis* species complex *E. carolleeae* (Atlantic clade), *E. gulfia* (Gulf clade), and *E. affinis* proper (Europe clade), their sequences were manually obtained by performing local BLAST of their reference genomes (Du et al., 2025). For certain taxonomic groups with limited genomic resources, αCA sequences were mined from transcriptome sequences (Supplementary Material; Table S1). Only BLAST query hits with E-value < 10^-5^were added to the initial dataset. When multiple alternative splice variants were present, the sequences with the longest Open Reading Frame (ORF) were retained. For quality control, all paralogs were screened for completeness. Incomplete paralogs were marked and removed from subsequent analyses. After the quality control processes, a total of 294 arthropod αCA sequences, out of the initial 351, were retained as complete and non-partial. From the initial 351 arthropod αCA sequences, the number of non-CARP αCA sequences and CARP αCA sequences were summarized for each arthropod order and subphylum (Supplementary Material; Table S7).

### 4.2 *In Silico* Subcellular Localization Prediction

The subcellular localization of complete amino acid sequences of arthropod and chordate αCA paralogs was predicted through the software DeepLoc 2.0 (Thumuluri et al. 2022). DeepLoc 2.0 takes amino acid sequences of eukaryotic proteins to predict the protein subcellular localization through a multi-staged deep learning approach. The DeepLoc tool is trained through the UniProt database (UniProt Consortium, 2019). DeepLoc is also capable of predicting multiple localizations for a given protein sequence, delineated with a “/”. For example, a predicted localization of “Extracellular/Golgi Apparatus” indicates that the protein is predicted to localize both extracellularly and on Golgi.

The proportion of αCA homologs belonging to each distinct αCA clade (Fig. 3a, 4a-b, Supplementary Material; Fig. S8) and predicted subcellular localization category (Supplementary Material; Fig. S7) was determined for arthropod and chordate αCA. For this analysis, species with less than or equal to 5 non-CARP αCA paralogs were excluded, such as *Aphonopelma hentzi, Calanus finmarchicus, Centruroides vittatus, Eupolybothrus cavernicolus, Stylopallene cheilorhynchus, Tachypleus tridentatus, Tanystylum orbiculare*, *Adelges cooleyi*, *Aricia agestis*, and *Ciona intestinalis*. The purpose of this exclusion was to reduce biases in proportion, due to the lower value in the denominator (the total number of non-CARP αCA paralogs). For the assessment of proportion of αCA homologs belonging to each predicted subcellular localization category (Supplementary Material; Fig. S7), CARP sequences were removed. The purpose of this exclusion is to reduce biases from the CARP expansion seen in decapods, which would skew the proportion of extracellularly localized αCA homologs in arthropods.

### 4.3 Phylogenetic Reconstruction

A comprehensive αCA phylogeny was reconstructed with 298 (294+4) of amino acid sequences from 31 arthropod taxa and 1 outgroup species (See Data Availability section). The 294 complete αCA amino acid sequences of each arthropod taxa, in addition to the 4 Porifera αCA paralogs, were aligned using the default FFT-NS-2 algorithm implemented in MAFFT Version 7.5 (Katoh et al. 2013). The resulting alignment was then trimmed using ClipKIT v.2.2.6 with the “kpic-smart-gap” algorithm, which retains only parsimony informative sites and constant sites, while removing all sites with “gappyness” above a dynamically determined threshold (0.9933% gap for this given alignment) (Steenwyk et al. 2020).

The trimmed alignment was then used to reconstruct a Maximum Likelihood phylogeny using IQTree v.2.2.0.3, with 100,000 ultrafast bootstrap iterations and the Q.pfam+R8 model of evolution (Minh et al. 2020; Minh et al. 2021). The evolutionary model with the best fit to the data was determined based on BIC through the ModelFinder Plus (-MPF) algorithm implemented within the IQTree software suit (Kalyaanamoorthy et al. 2017). The tree was rooted on the ancestral node of the 4 Porifera αCA paralogs. Graphical visualization, including the mapping of the predicted subcellular localization, was performed with TreeViewer (Bianchini et al. 2024).

To determine signatures of selection (based on *dN/dS*) through a phylogenetic approach using HyPhy (Pond et al. 2005) (see section 4.5 below), we reconstructed three additional arthropod αCA phylogenies based on the three distinct arthropod αCA clades (Extracellular & Membrane-bound, Cytosolic, and CARP clade) that we defined based on the comprehensive arthropod phylogeny of 298 sequences (see above). These phylogenies were reconstructed using the exact same procedure as the comprehensive arthropod αCA phylogeny discussed above (with 294+4 sequences). The Bayesian information criterion (BIC) best fit model for the Extracellular & Membrane-bound, Cytosolic, and CARP clade phylogenies were selected as Q.pfam+R8, LG+I+G4, and LG+I+I+R5, respectively, through the ModelFinder Plus algorithm (Kalyaanamoorthy et al. 2017). These phylogenies were unrooted and visualized with TreeViewer (Bianchini et al. 2024).

A phylogeny of 72 chordate αCA paralogs was also reconstructed using the same procedures described above. The comprehensive chordate αCA phylogeny was rooted on the ancestral node of the same 4 Porifera αCA paralogs. Through the ModelFinder Plus algorithm, the BIC best fit model for the comprehensive chordate αCA phylogeny was determined as Q.pfam+I+I+R4 (Kalyaanamoorthy et al. 2017). To compare the phylogenetic topology and predicted subcellular localization between the chordate & arthropod αCA, three additional clade-specific chordate αCA phylogenies (Extracellular & Membrane-bound, Cytosolic, and CARP clade) were generated using the same procedure as for arthropods. The BIC best fit model for the Extracellular & Membrane-bound, Cytosolic, and CARP clade chordate αCA phylogenies were Q.pfam+I+I+R3, LG+I+G4, JTT+G4, respectively, using the ModelFinder Plus algorithm (Kalyaanamoorthy et al. 2017).

### 4.4 Alignment-free αCA Sequence Clustering

To provide further support to the topology of our arthropod αCA phylogeny, we constructed a UMAP projection of the arthropod αCA, using an alignment-free method (Yeung et al. 2023). Under their scripts, the ESM-1b protein language model was applied to unaligned amino acid sequences of arthropod αCA paralogs (Rives, Meier, Sercu et al., 2021). Large protein language models, such as the ESM-1b, are trained on a vast number of publicly available protein sequences. By applying this language model to a given set of protein sequences, it generates various numerical representations of the protein sequence, or sequence embedding, which elucidates the biochemical properties and structural features of each protein. These sequence embeddings are mapped as a high-dimensional vector. The distance between the representations of each of the protein sequences is calculated through a range of distance metrics, including cosine distance and Triangle Similarity-Sector Similarity (TS-SS). The distance matrix is subjected to further dimensionality reduction via UMAP, creating a two-dimensional projection with uniform manifold approximation for easier visualization (McInnes et al. 2018).

Unedited amino acid sequences of arthropod αCA were parsed through the scripts described in Yeung et al., (2023) (https://github.com/esbgkannan/chumby) (See Github Repository).

### 4.5 Signatures of Selection along Phylogenetic Branches and at Amino Acid Sites

Signatures of selection along phylogenetic branches and at amino acid sites were examined in the clade-specific arthropod αCA phylogenies using the software HyPhy (Pond et al. 2005). The signatures of selection here were quantified through the ω value (the ratio of synonymous to nonsynonymous substitution rates; *dN/dS*) as implemented in the HyPhy batch language.

To compute and compare the mean *dN/dS* among the arthropod αCA clades, among the chordate αCA clades, and between arthropod and chordate αCA clades, untrimmed amino acid alignments that included paralogs from chordate and arthropod αCA clades were generated. To ensure homology while reducing saturation, any amino acid residue with less than 50% homology was trimmed using the Custom Site Trimming (cst) mode implemented in ClipKIT v.2.2.6 (Steenwyk et al. 2020). The codons with less than 50% homology were determined using a custom python script (see github repository; cst_alignment_homology_getter.py).

The resulting trimmed amino acid alignments were converted to codon alignments through the PAL2NAL web server (https://www.bork.embl.de/pal2nal/) (Suyama et al. 2006). The PAL2NAL program takes an amino acid alignment and its corresponding nucleotide sequence as an input and transforms the nucleotide sequence into a codon alignment based on the amino acid alignment. To ensure that the nucleotide sites that were compared between the different αCA clades were homologous, the αCA paralogs from different αCA clades were aligned to each other and filtered simultaneously. For example, when comparing between the arthropod and chordate CARP αCA clades, the CARP paralogs of the arthropod and chordate species were aligned and filtered together, so that there was a homologous residue in the edited chordate CARP sequence for each residue within the edited arthropod CARP sequence.

Branch and node-specific ω values (*dN/dS*) were then calculated through FitMG94 for each arthropod and chordate αCA clades, taking the translated codon alignments and clade-specific unrooted phylogeny as inputs (Pond et al. 2005). The FitMG94 analysis fits a MG94 codon substitution model on an alignment and a tree to report the expected synonymous and non-synonymous substitutions per nucleotide side for each branch and node of the input (Muse & Gaut, 1994). To further eliminate saturation, any branches and nodes with a synonymous substitution rate greater than 3 or less than 0.0001 (*d_S_* >3, *d_S_* <0.0001) or with ω values greater than 3 (*dN/dS*> 3) were removed. The resulting ω values (*dN/dS*) were visualized using Raincloud plots (Allen et al. 2019).

To investigate the signatures of positive selection along each phylogenetic branch, the Adaptive Branch-Site Random Effects Likelihood (aBSREL) method in the software package HyPhy was utilized (Suyama et al. 2006; Smith et al. 2015; Pond et al. 2005; Weaver et al., 2018). As a first step, this analysis used untrimmed amino acid alignments for each arthropod αCA clade. To eliminate saturation due to misalignment, any amino acid residues with less than 60% homology were trimmed using the Custom Site Trimming (cst) mode implemented in ClipKIT v.2.2.6 (Steenwyk et al. 2020). Such trimming removes codon sites with differing strengths of negative selection between sequences, a known confounding factor in the detection of positive selection (Chen et al., 2022). This analysis allows us to focus our selection analysis on potentially functional sites that have high sequence homology between species. Note that the homology threshold used for this selection analysis was more stringent than that used to compute the *dN/dS*. The trimmed clade-specific arthropod αCA alignments were converted to codon alignments through the PAL2NAL webserver (https://www.bork.embl.de/pal2nal/) (Suyama et al. 2006). Using the previously constructed phylogeny (see section 4.3), branch-specific signatures of positive selection were investigated using aBSREL (Suyama et al. 2006; Smith et al. 2015; Pond et al. 2005; Weaver et al., 2018). Each arthropod αCA clade was tested independently for signatures of positive selection.

Using aBSREL, branches leading each αCA paralog of the *Eurytemora affinis* species complex were selected as the foreground. The purpose was to test the hypothesis that αCA paralogs of *E. affinis* complex previously shown to have signatures of selection associated with contemporary salinity change (Lee, 2021; Stern and Lee, 2020; Stern et al., 2022; Diaz et al., 2023), also show signatures of selection based on *dN/dS* in a comparison with arthropod clades. In this analysis, we selected each αCA paralog of the *Eurytemora affinis* species complex as the foreground, and compared them to the general arthropod αCA background. As an exploratory analysis, we also set all arthropod αCA branches as foreground to elucidate any non-*Eurytemora* αCA that show signatures of natural selection. A Holm-Bonferroni corrected *P*-value of 0.05 was used as a threshold for significance (Holm,1979; Smith et al. 2015).

In addition, Extracellular & Membrane-bound αCA sequences of *E. affinis* complex that showed signatures of positive selection in aBSREL were scanned for site-specific signatures of positive selection using Branch-Site Unrestricted Statistical Test for Episodic Diversification (BUSTED) and Mixed Effect Model of Evolution (MEME) in HyPhy (Suyama et al. 2006; Murrell et al. 2012; Murrell et al., 2015; Pond et al. 2005; Weaver et al., 2018). For BUSTED, we selected the basal node of *E. carolleeae* and *E. gulfia* αCA12 (Node 7) and its terminal branches as the foreground for testing and the general arthropod Extracellular & Membrane-bound αCA homologs as the background. We interpreted the Evidence Ratio (ER) values generated from BUSTED as an *exploratory tool* to understand which nucleotide sites from αCA12 contributed to the signals of positive selection seen in BUSTED and aBSREL. Note that ER provides information on whether a nucleotide site could evolve under positive selection and contribute to the gene-wide signal of positive selection. However, this parameter does not provide any statistical evidence that these nucleotide sites are under positive selection (Murrell et al., 2015). Therefore, this result remains strongly exploratory. The significance threshold of ER was determined using an evidence ratio threshold of 10. For MEME, we implemented a *P*-value of 0.05 as the threshold for significance.

### 4.6 Protein structure modeling

A 3D protein structure of αCA paralog 12 (αCA12) of *Eurytemora carolleeae* was modeled using AlphaFold 3 (Abramson et al. 2024) (https://alphafoldserver.com/welcome). The resulting protein model was visualized through the PyMol software suit, where nucleotide sites under positive selection were highlighted (DeLano 2002). Confidence of the folded αCA12 protein model is available through the plDDT metric (Supplementary Material; Fig. S9).

## 5. Data and Resource Availability

Data, including raw sequences, some intermediary files, and final edited sequence to reproduce the analyses can be found online in the supplementary materials and at GitHub repository (https://github.com/Yf-Joye-Z/arthropod_aCA).

## Acknowledgements

This work was supported by the University of Wisconsin-Madison Integrative Biology Summer Research Fellowship, Sophomore Research Fellowship, Hilldale Research Fellowship, and Genetics and Genomics Undergraduate Distinguished Research Fellowship to Yifei Joye Zhou, and National Science Foundation grants IOS-2412790, OCE-1658517, and DEB-2055356, and French National Research Agency ANR-19-MPGA-0004 (Macron’s “Make Our Planet Great Again” award) to Carol E. Lee. Benjamin Wilson and Cullen Meyer shared their αCA sequence data. Alexander C. Taylor provided training on general bioinformatics and software usage, and reviewed the manuscript. Lee Lab members Zhengyong Du, Patricia Zito, Jinhui Wang, and Caitlin McDonough-Goldstein provided research insights, scientific discussions, and training on the concepts and methodologies involved in this project.

