## supplementary_figures for "Evolutionary history of the *α-Carbonic Anhydrase* (*αCA*) gene family in the phylum Arthropoda, with a focus on the copepod *Eurytemora affinis* species complex"

### Supplementary Material

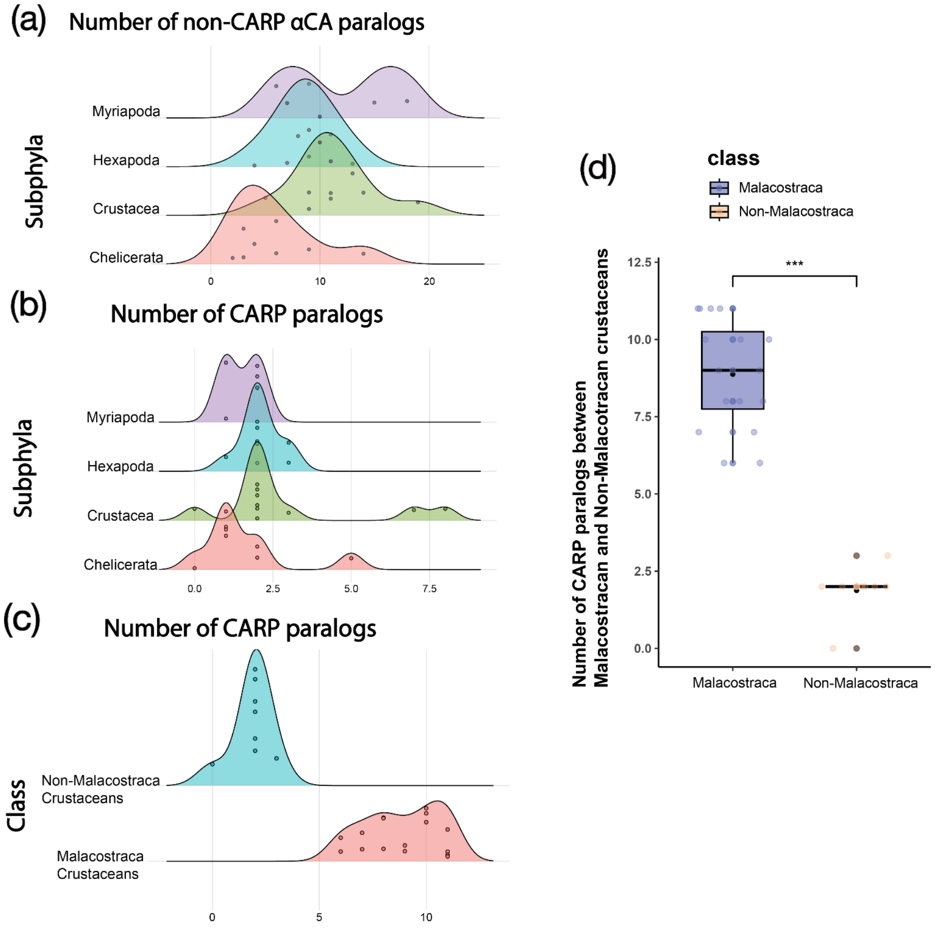


**Supplementary Figure 1. Distribution of CARP and non-CARP αCA from different Arthropod subphyla. (a)** Ridge density plot showing the number of CARP paralogs of species from the four arthropod subphyla. (**b**) Ridge density plot showing the number of non-CARP αCA paralogs of species from the four arthropod subphyla. (**c**) Ridge density plot showing the number of CARP paralogs of species from Malacostracan and Non-Malacostracan crustaceans. Note that Malacostracan crustaceans have an extraordinarily high CARP paralog count when compared to the Non-Malacostraca crustaceans. (**d**) Proportion of CARP paralogs per species between Malacostracan and Non-Malacostracan crustaceans. W=128. ns: not significant; * = p<0.05; ** = p<0.01; *** = p<0.001; **** = p<0.0001; Wilcoxon rank-sum test. .

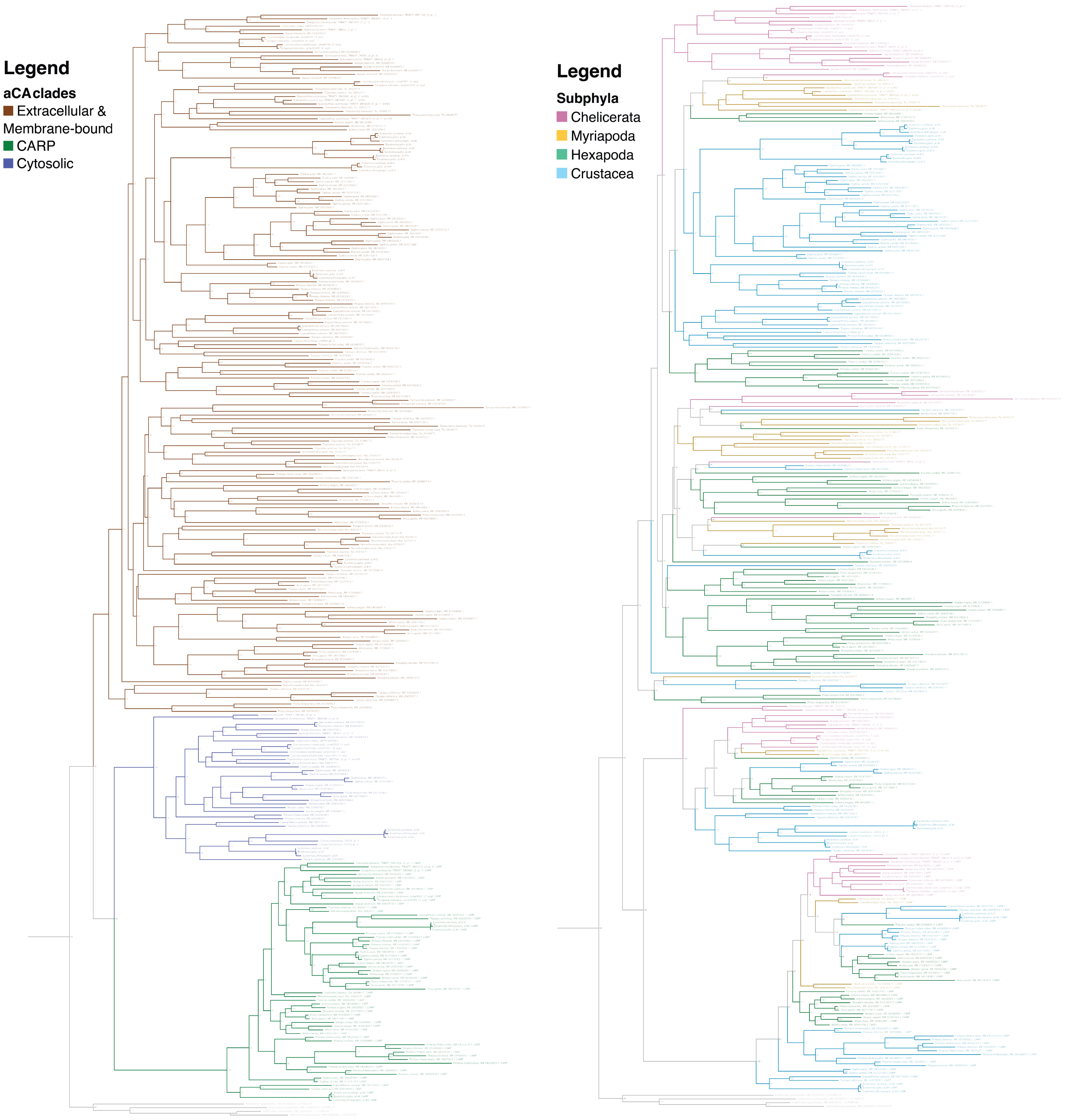


**Supplementary Figure 2. Phylogeny of arthropod αCA .** Maximum-Likelihood gene tree of arthropod αCA, inferred from IQ-tree (Minh et al., 2020). The phylogeny corresponds to Figure 1 and 2 of the main text. The phylogeny on the left is color coded based on the three distinct αCA clades. The phylogeny on the right is color coded based on the subphyla of the species from which the paralog was mined.

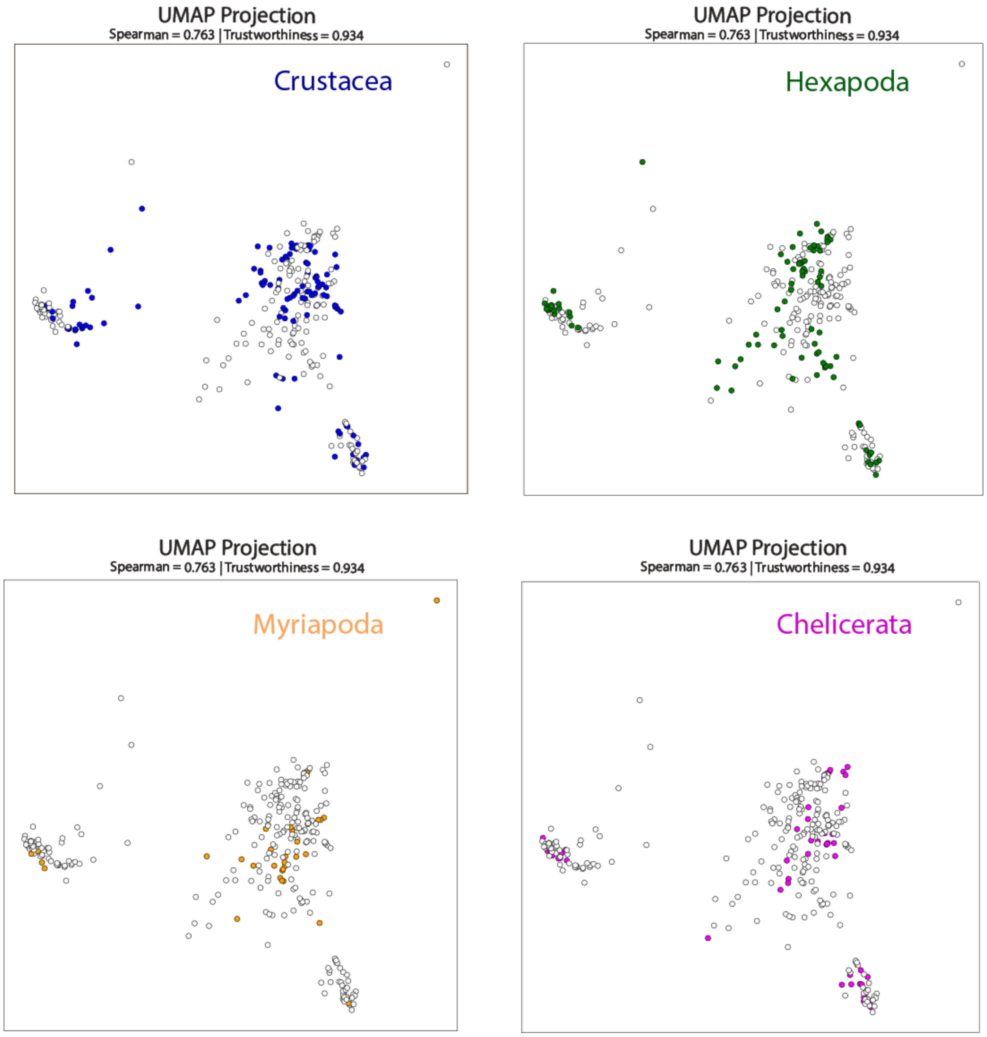


**Supplementary Figure 3. Distribution of αCA from four arthropod subphyla across the 3 arthropod αCA clades.** The αCA homologs from each arthropod subphylum separated into the three arthropod αCA clades (Extracellular & Membrane-bound, Cytosolic, CARP). UMAP projection of 294 distinct arthropod αCA homologs based on ESM-1b embedding, similar to Figure 1b & 2b. The distance was calculated through TS-SS similarity, and the global and local accuracy of this clustering is shown through a Spearman rank correlation and trustworthiness value. Each dot represents a distinct αCA paralog. The colored dots in each plot represents the αCA orthologs & paralogs for each respective subphyla, Crustacea, Hexapoda, Myriapoda, and Chelicerata.

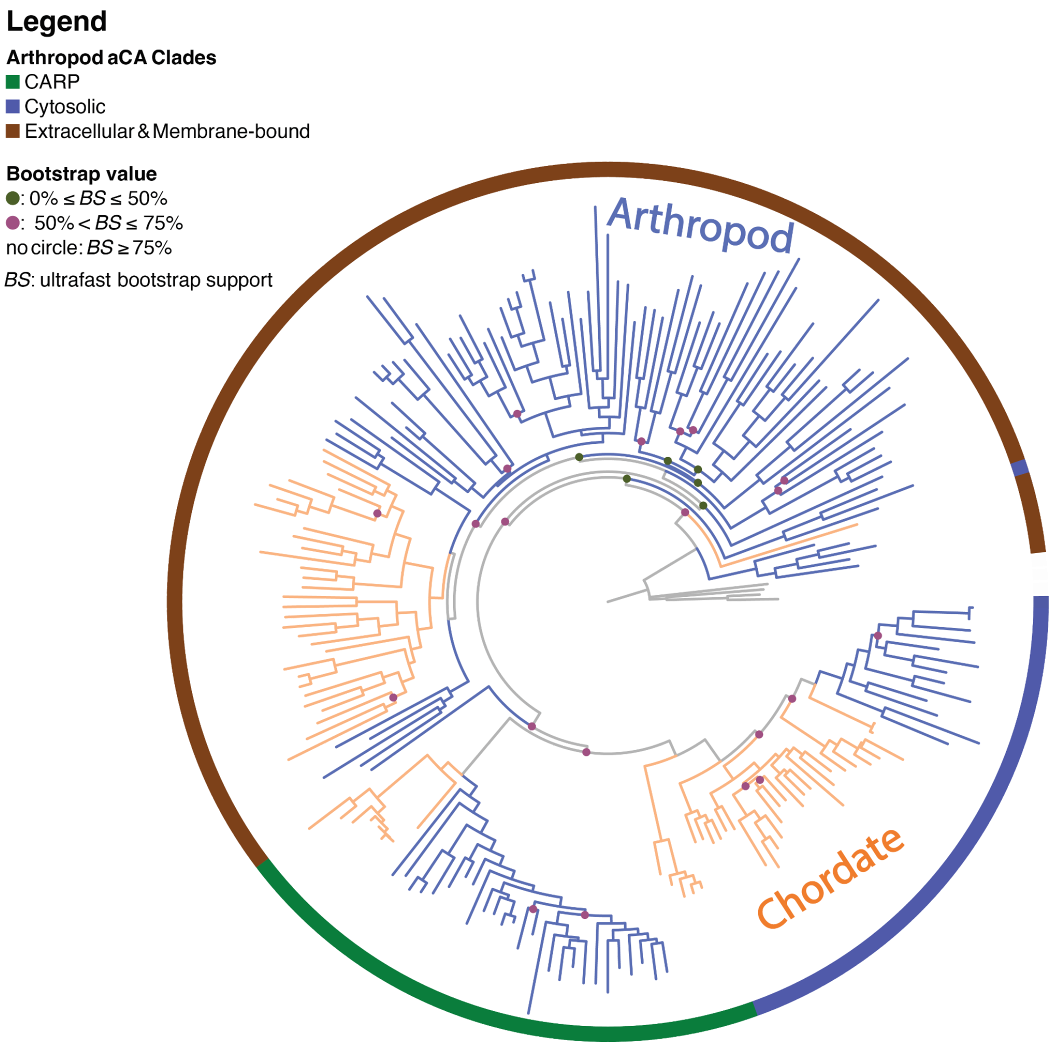


**Supplementary Figure 4. Phylogeny of arthropod and chordate αCA.** Maximum-Likelihood gene tree of arthropod and chordate αCA, inferred from IQ-tree (Minh et al., 2020). The phylum identity of each branch is coded by its color, with blue corresponding to arthropods and yellow corresponding to chordates. The outer circle denotes the 3 distinct αCA clades.

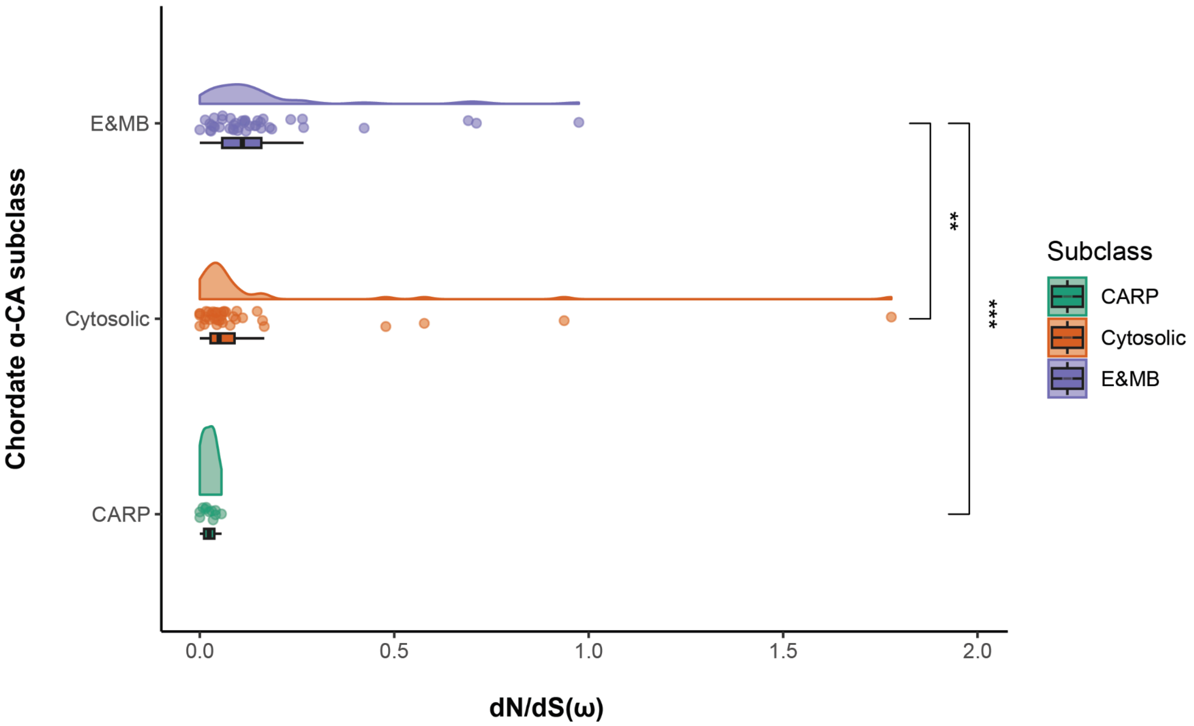


**Supplementary Figure 5. Chordate Extracellular & Membrane-bound αCA show similar patterns of mean *dN/dS* as arthropod αCA.** Box and density plots showing the branch and node ω value (*dN/dS*) of each chordate αCA clade. ω values (*dN/dS*) were obtained by fitting the MG94 model onto the edited chordate αCA sequences (see Methods and Materials). E&Mb stands for the Extracellular & Membrane-bound αCA clade. E&Mb: mean ω = 0.16 ± 0.03 SE. Cytosolic: mean ω = 0.14 ± 0.05 SE. CARP: mean ω = 0.02 ± 0.005 SE. E&Mb vs. Cytosolic: W=502, E&Mb vs. CARP: W=48. ns = not significant; * = p<0.05; ** = p<0.01; *** = p<0.001; **** = p<0.0001; Wilcoxon rank-sum test.

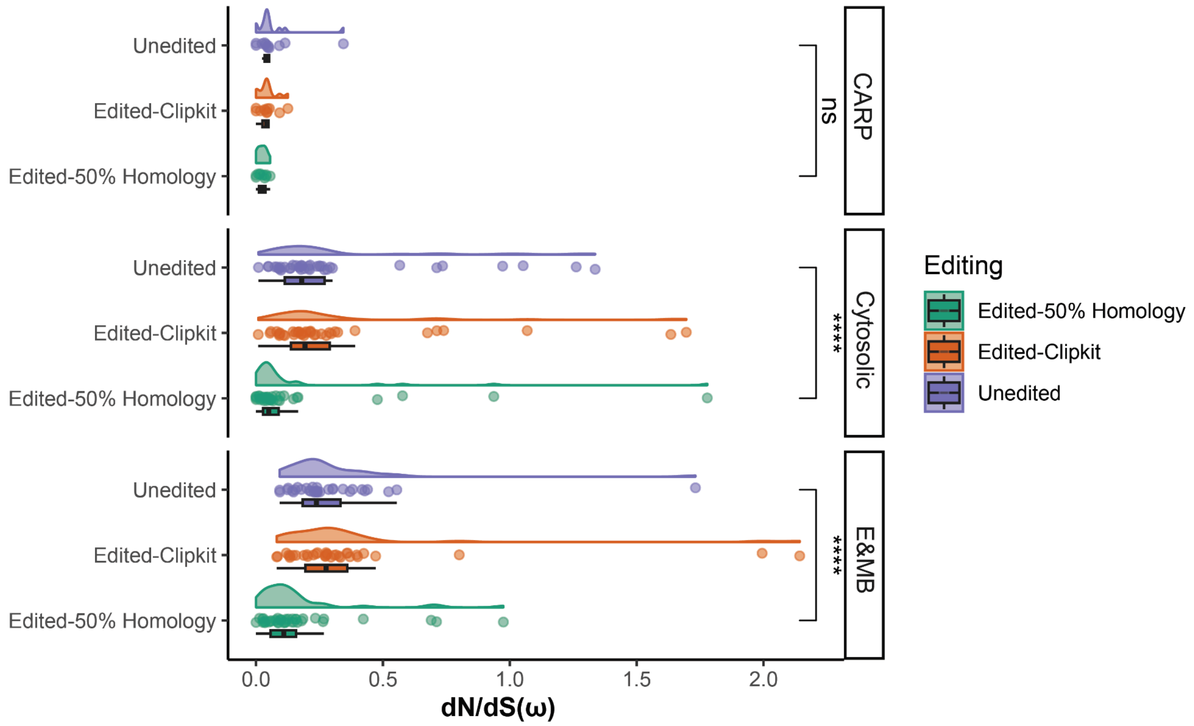


**Supplementary Figure 6. Comparison of ω values (*dN/dS*) from Chordate αCA using different alignment editing methods.** Box and density plots showing the branch and node-specific ω value (*dN/dS*) of each chordate αCA clade, compared between alignments that are unedited, edited using kpic-smartgap as implemented in ClipKit, and edited by removing any codons that have less than 50% homology across the alignment. E&Mb stands for the Extracellular & Membrane-bound αCA clade. Unedited vs. Edited - 50% homology for CARP: W=40, Cytosolic: W=264, E&Mb: W=259. ns = not significant; * = p<0.05; ** = p<0.01; *** = p<0.001; **** = p<0.0001; Wilcoxon rank-sum test

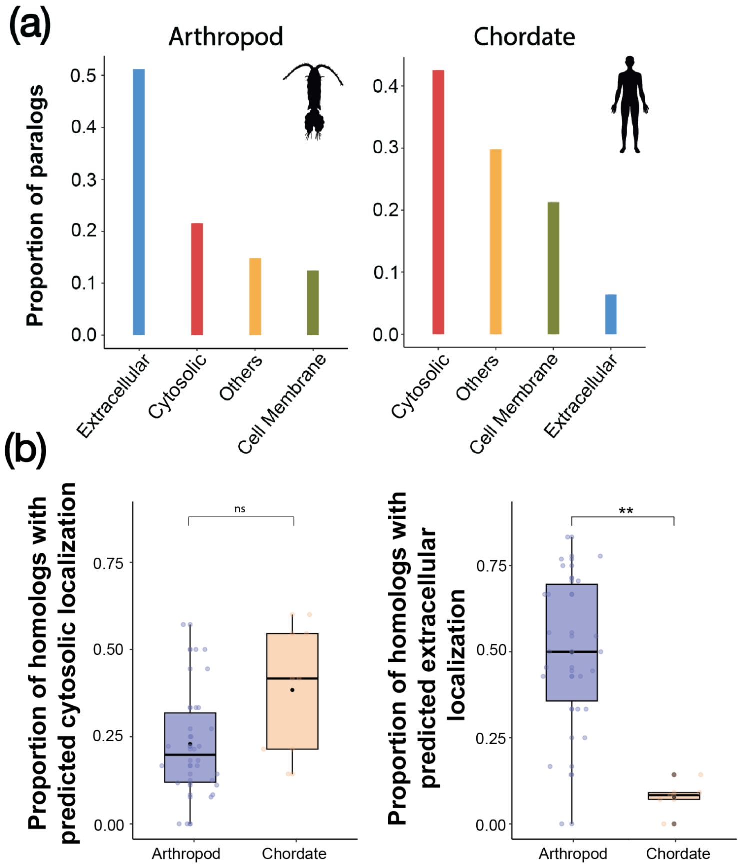


**Supplementary Figure 7. Differential αCA gene family expansion patterns in arthropods versus chordates** (**a**) Proportion of αCA homologs (orthologs and paralogs) with predicted subcellular localization of Extracellular Cytosolic, Cell Membrane, or Others over the total number of arthropod αCA homologs (left) and chordate αCA homologs (right). (**b**) Boxplots comparing the proportion of homologs with predicted cytosolic localization (left) and predicted extracellular localization (right) per species between arthropod and chordate species. Cytosolic: Arthropod mean = 0.23 ± 0.03 SE, Chordate mean = 0.38 ± 0.09 SE. E&Mb: Arthropod: mean = 0.50 ± 0.05 SE, Chordate mean = 0.08 ± 0.02 SE. Wilcoxon rank-sum test, ns: not significant; * = *P* < 0.05; ** = *P* < 0.01. Cytosolic Arthropod vs. Chordate: W=30. Extracellular Arthropod vs. Chordate: W=105.

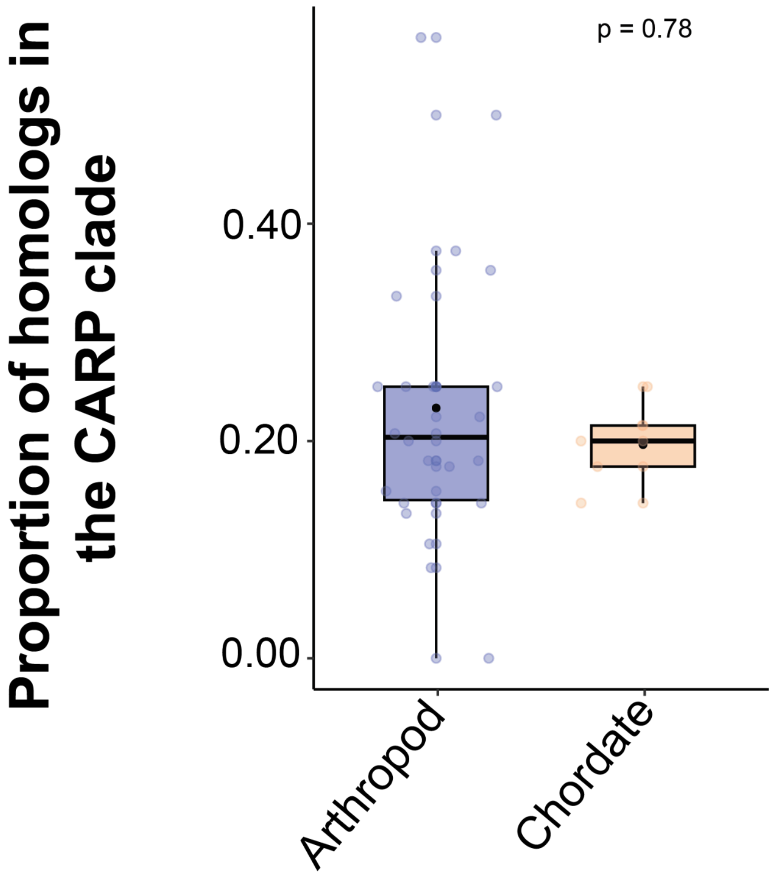


**Supplementary Figure 8. Proportion of αCA homologs within the CARP clade in arthropods and chordates**. Proportion of homologs per species that are within the CARP clade, one of the 3 distinct αCA clades described in this study. Wilcoxon rank-sum test, W = 60, *P* = 0.78. Arthropod mean = 0.23 ± 0.03 SE. Chordate mean = 0.20 ± 0.02 SE.

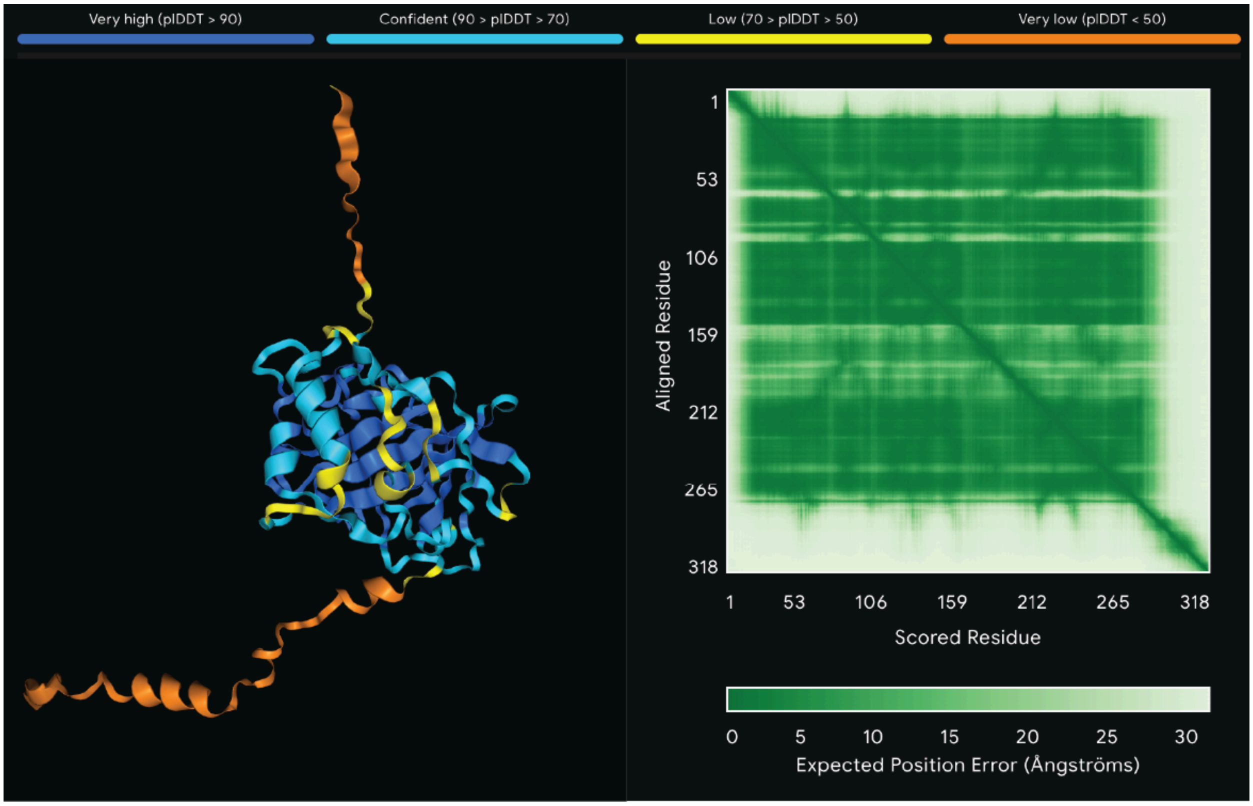


**Supplementary Figure 9. Structural Confidence of the folded *Eurytemora carolleeae* αCA12 protein.** Confidence metrics and expected per-residue position error matrix generated from the AlphaFold3 server. plDDT = per-atom confidence estimate, where higher values mean higher confidence.

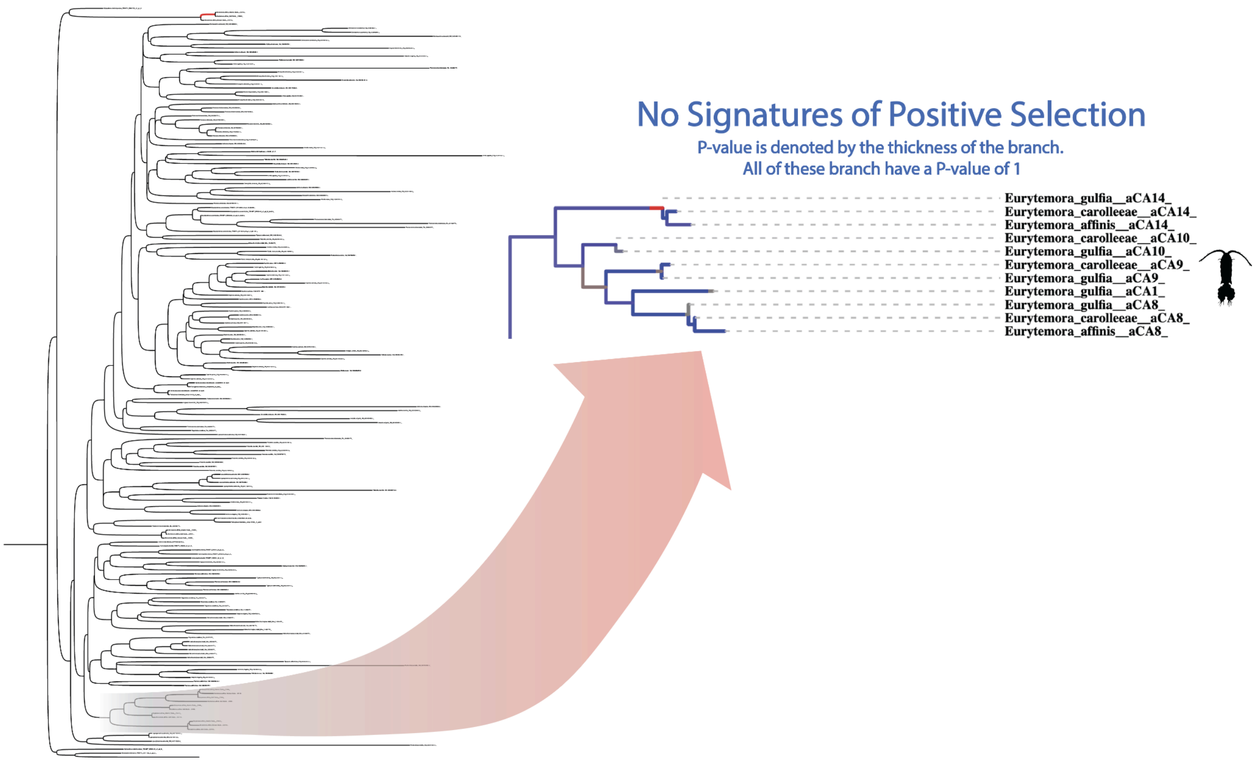


**Supplementary Figure 10. αCA9 of *Eurytemora affinis* species complex shows no significant signatures of positive selection.** Extracellular & Membrane-bound arthropod αCA phylogeny. The highlighted branch is expanded on the right. Corrected P-value that corresponds to the signatures of positive selection is denoted by the thickness of each branch. The signatures of positive selection are determined through the HyPhy software, aBSREL (Smith et al., 2014). None of the highlighted *E. affinis* αCA paralogs show signatures of natural selection. The image is taken from HyPhy Vision (http://vision.hyphy.org/).

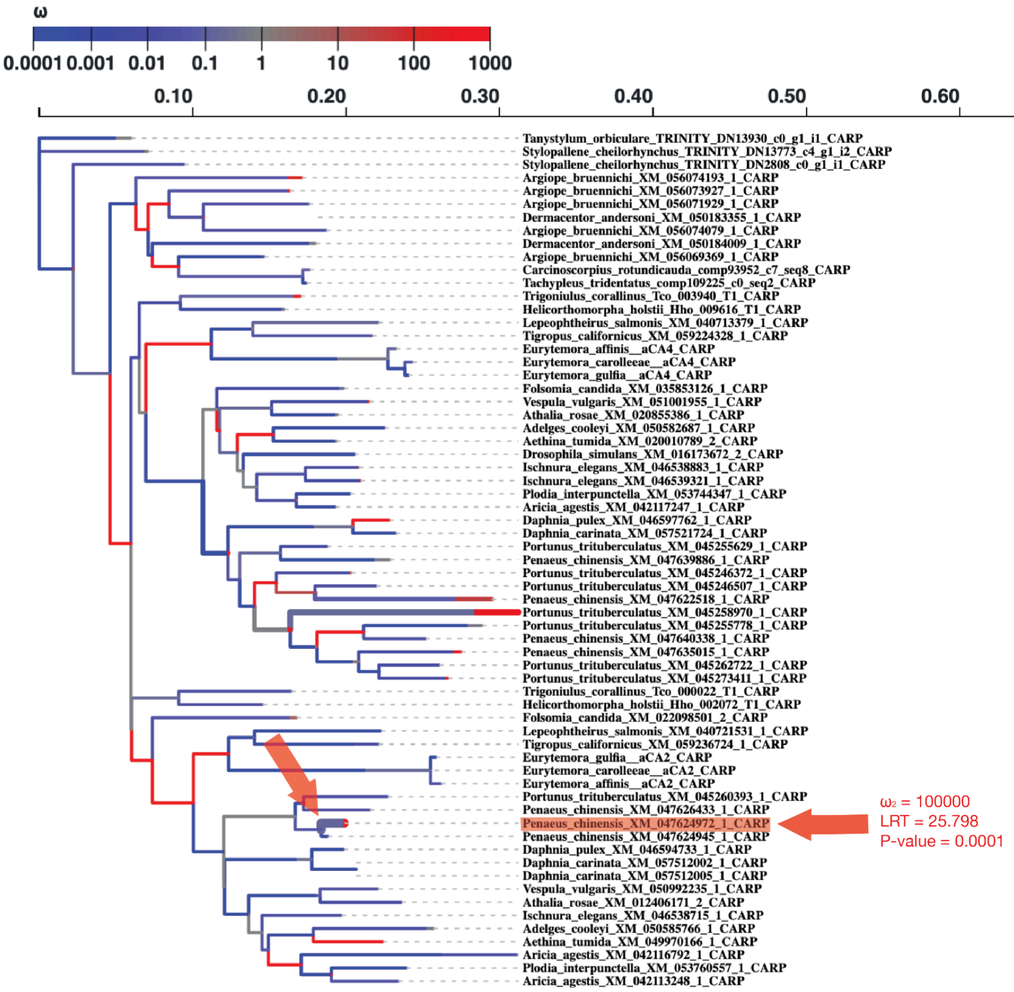


**Supplementary Figure 11. Branch-level signatures of positive selection within the arthropod CARP phylogeny.** The colors of the phylogenetic branches correspond to the ω values of the branches, with red representing higher values and blue representing lower values. The significance of signatures of positive selection is denoted by the thickness of each branch. The signatures of positive selection are determined through the HyPhy software, aBSREL (Smith et al., 2014). The one branch/node with signatures of positive selection is highlighted using an arrow, with the associated w, LRT, and P-values shown next to the arrows on the right. The image is taken from HyPhy Vision (http://vision.hyphy.org/).

### Supplementary Methods

#### 1.1 Gene mining for Malacostracan αCA sequences

To gain further insights into the high number of CARPs among the 2 decapod species (*Penaeus chinensis*, *Portunus trituberculatus*), we mined CARP paralogs from 13 additional decapod species and 1 amphipod species (Supplementary Table 8). Using the CARP sequences from *Penaeus chinensis* as a query, BLAST searches (blastp) were conducted on each of these genomes (Altschul et al., 1990; Camacho et al., 2009; Johnson et al., 2008). Only BLAST query hits with E-value < $10^{-5}$ were added to the initial dataset. When multiple alternative splice variants were present, the sequences with the longest Open Reading Frame (ORF) were retained.

**1.2 Phylogenetic reconstruction of Arthropod and Chordate αCA**

To unravel the grouping pattern of arthropod and chordate αCA within each of the distinct αCA phylogenetic clades, we reconstructed a Maximum-Likelihood phylogeny of αCA using sequences from both arthropods and chordates. To be specific, this phylogeny included αCA sequences from 10 species of arthropods (*Dermacentor andersoni, Argiope bruennichi, Thereuonema tuberculata, Helicorthomorpha holstii, Eurytemora carolleeae, Portunus trituberculatus, Daphnia pulex, Ischnura elegans, Plodia interpunctella, Drosophila simulans*) and 6 species of chordates (*Homo sapiens, Danio rerio, Xenopus tropicalis, Gallus gallus, Ciona intestinalis, Branchiostoma lanceolatum*). The complete αCA amino acid sequences of these arthropod and chordate species, in addition to the 4 αCA paralogs from the outgroup Porifera species, were aligned using the default FFT-NS-2 algorithm implemented in MAFFT Version 7.5 (Katoh et al. 2013). Amino acid sequences were used rather than nucleotide sequences. The resulting alignment was then trimmed using ClipKIT v.2.2.6 with the "kpic-smart-gap" algorithm, which retains only parsimony informative sites and constant sites, while removing all sites with "gappyness" above a dynamically determined threshold (0.9949% gap for this given alignment) (Steenwyk et al. 2020).

The trimmed alignment was then used to reconstruct a Maximum Likelihood phylogeny using IQTree v.2.2.0.3 (reference), with 100,000 ultrafast bootstrap iterations and the Q.pfam+R10 model of evolution (Minh et al. 2020; Minh et al. 2021). The evolutionary model with the best fit to the data was determined based on BIC through the ModelFinder Plus (-MPF) algorithm implemented within the IQTree software suit (Kalyaanamoorthy et al. 2017). The phylogeny was rooted on the ancestral node of the 4 Porifera αCA paralogs. Graphical visualization, including the mapping of the predicted subcellular localization, was performed with TreeViewer (Bianchini et al. 2024).
